# Generation of comprehensive ecosystems-specific reference databases with species-level resolution by high-throughput full-length 16S rRNA gene sequencing and automated taxonomy assignment (AutoTax)

**DOI:** 10.1101/672873

**Authors:** Morten Simonsen Dueholm, Kasper Skytte Andersen, Simon Jon McIlroy, Jannie Munk Kristensen, Erika Yashiro, Søren Michael Karst, Mads Albertsen, Per Halkjær Nielsen

**Affiliations:** Center for Microbial Communities, Department of Chemistry and Bioscience, Aalborg University, Aalborg, Denmark

## Abstract

High-throughput 16S rRNA gene amplicon sequencing is an essential method for studying the diversity and dynamics of microbial communities. However, this method is presently hampered by the lack of high-identity reference sequences for many environmental microbes in the public 16S rRNA gene reference databases, and by the absence of a systematic and comprehensive taxonomy for the uncultured majority. Here we demonstrate how high-throughput synthetic long-read sequencing can be applied to create ecosystem-specific full-length 16S rRNA gene amplicon sequence variant (FL-ASV) reference databases that include high-identity references (>98.7% identity) for nearly all abundant bacteria (>0.01% relative abundance) using Danish wastewater treatment systems and anaerobic digesters as an example. In addition, we introduce a novel sequence identity-based approach for automated taxonomy assignment (AutoTax) that provides a complete seven-rank taxonomy for all reference sequences, using the SILVA taxonomy as a backbone, with stable placeholder names for unclassified taxa. The FL-ASVs are perfectly suited for the evaluation of taxonomic resolution and bias associated with primers commonly used for amplicon sequencing, allowing researchers to choose those that are ideal for their ecosystem. The AutoTax taxonomy greatly improves the classification of short-read 16S rRNA gene amplicon sequence variants (ASVs) at the genus- and species-level, compared to the commonly used universal reference databases. Importantly, the placeholder names provide a way to explore the unclassified environmental taxa at different taxonomic ranks, which in combination with *in situ* analyses can be used to uncover their ecological roles.

## Introduction

Microbial communities underpin key biochemical transformations in natural and engineered ecosystems. A deep understanding of these systems requires reliable identification of the microbes present, which can then be associated with their metabolic and ecological functions. Identification at the lowest taxonomic rank is preferred, as microbial traits vary in their degree of phylogenetic conservation, and many ecologically important traits are conserved only at the genus and species level (1).

Identification of microbes is commonly achieved by high-throughput 16S rRNA gene amplicon sequencing, where a segment of the 16S rRNA gene spanning one to three hypervariable regions (usually 200-500 bp long) is amplified by PCR and sequenced. The amplicons are then clustered into operational taxonomic units (OTUs) or used to infer exact amplicon sequence variants (ASVs) with denoising algorithms such as Deblur (2), DADA2 (3), or Unoise3 (4). The taxonomy of these amplicons is then assigned based on their comparison to a reference database (5–7).

ASVs are often preferred over OTUs because they provide single-nucleotide resolution and can be applied as consistent labels for microbial identification independently of a 16S rRNA gene reference database (8). This approach is used with short-read ASVs in several large-scale projects, including the Earth Microbiome Project (EMP) (9) and the American Gut project (10), to provide detailed insight into the factors that shape the overall microbial community diversity and dynamics. However, ASVs are not ideal as references for linking microbial identity with the physiology and ecology of key community members, which is crucial if we want to use microbial community structure to predict ecosystem functions or process performance in engineered systems. Firstly, without taxonomic assignment, it is not possible to compare results across studies that have used primers targeting different regions of the 16S rRNA gene. Secondly, short-read ASVs alone do not contain enough evolutionary information to resolve their phylogeny confidently (11, 12). This makes it impossible to report and infer how microbial traits are conserved at different phylogenetic scales. As such, functional properties for uncultured lineages predicted from the annotation of metagenome-assembled genomes (MAGs) with complete rRNA genes (Hiqh-quality MIMAGs standard) (11), or determined through *in situ* studies, cannot be confidently linked to the shorter ASV sequences (12, 13). A robust taxonomic assignment is therefore crucial for cross-study comparisons and the accumulation and dissemination of knowledge relating to uncultured lineages.

Taxonomic assignment to ASVs relies on a classifier, e.g., the SINTAX (7) or the RDP classifier (6), which uses statistical algorithms to compare each ASV to a full-length 16S rRNA gene reference database to propose the best estimate of their taxonomy. Confident classification at the lowest taxonomic ranks (genus and species) requires high-identity reference sequences (≥98.7% identity) and a complete seven-rank taxonomy for all references (13). Neither of these criteria is met with the most commonly applied universal reference databases, e.g., Greengenes (14), SILVA (15), and RDP (16), as they lack sequences or taxonomic assignment for many uncultivated environmental taxa. Given the vast diversity predicted for microbial life on Earth (17), it will be some time before reference sequences for all species are generated, and the manual curation of their taxonomy will not be feasible.

A potential solution to the problems mentioned above is to create ecosystem-specific reference databases. These can either be ecosystem-curated versions of universal databases or smaller independent databases that only include sequences from the specific ecosystem. The MiDAS 2 database for microbes in biological wastewater treatment systems (18) and the Dictyopteran gut microbiota reference Database (DictDb) (19) are examples of ecosystem-curated versions of the SILVA databases, where the taxonomies for the abundant and process-critical microorganisms are manually curated and maintained. However, the mostly manual taxonomic curation of central reference databases is time-consuming and subjective. The universal reference databases are also clustered at 99% identity or below to keep the unique sequences to a manageable number, which reduces the taxonomic resolution.

Examples of independent ecosystem-specific databases include the human intestinal tract 16S taxonomic database (HITdb) (20), the human oral microbiome database (HOMD) (21), the freshwater-specific FreshTrain database (22, 23), the honey bee gut microbiota database (24), and the rumen and intestinal methanogen database (25). While such databases have been shown to improve the rate of classifications for amplicons, they generally contain a relatively limited number of sequences and are therefore associated with an inherent risk of over- or misclassification if the sequence being classified is not represented in the database. To address this issue, Rohwer *et al.* introduced the TaxAss algorithm that classifies amplicons using two reference databases: a universal database and a small ecosystem-specific database (23). Amplicons are first mapped to the ecosystem-specific database to determine the percent identity with the best hit, and those above a user-defined threshold are classified using the ecosystem-specific database. The remaining sequences are classified using the more extensive universal database. While higher rates of classification were achieved, an issue with the approach is the potential for closely related sequences that fall on either side of the user-defined threshold, to receive very different taxonomies, especially if the ecosystem-specific database is not updated to reflect the evolving taxonomy of the universal reference database. As such, while current strategies to create ecosystem-specific databases have shown promise, there are critical issues that need to be resolved before their potential can be realized.

The recent development of methods for high-throughput full-length 16S rRNA gene sequencing, e.g., synthetic long-read sequencing on the Illumina platform (26, 27), along with PacBio (28) or Nanopore (29) consensus sequencing, now allows for the generation of millions of high-quality reference sequences within days. Importantly, these technologies now allow for the high-throughput generation of high-identity reference databases with broad coverage of the true diversity. However, improving sequence coverage alone will not solve the problem of poor taxonomic assignments for many uncultured taxa. We have therefore developed the AutoTax pipeline, which provides a simple and efficient strategy for the creation of comprehensive ecosystem-specific taxonomies that cover all seven taxonomic ranks. AutoTax uses the SILVA taxonomy as a backbone and provides stable placeholder names for unclassified taxa, based on *de novo* clustering of sequences according to statistically supported identity thresholds for each taxonomic rank (12). Importantly, AutoTax databases are easily updated with subsequent releases of the SILVA taxonomy – avoiding the divergence of generated ecosystem-specific taxonomies with the universal reference database. The strict computational nature of the taxonomy assignment means that it is objective and reproducible. The simplicity of the applied *de novo* clustering also ensures that the placeholder names are maintained even though the database is expanded with additional reference sequences.

We demonstrate the potential of the AutoTax method by sequencing almost a million full-length 16S rRNA gene sequences from Danish biological wastewater treatment and bioenergy systems. The sequences were denoised to resolve full-length exact amplicon sequence variants (FL-ASVs) with single-nucleotide resolution. Taxonomy was assigned to the FL-ASVs using the AutoTax pipeline to create an ecosystem-specific reference database. As evidence supporting the value of our approach, mapping of short-read amplicon data revealed that a substantially higher proportion of sequences were matched to high-identity references, and received species and genus level classification when the FL-ASV database was used compared to the much larger public universal reference databases.

## Materials and methods

### General molecular methods

Concentration and quality of nucleic acids were determined using a Qubit 3.0 fluorometer (Thermo Fisher Scientific) and an Agilent 2200 Tapestation (Agilent Technologies), respectively. Agencourt RNAClean XP and AMPure XP beads were used as described by the manufacturer, except for the washing steps, where 80% ethanol was used. RiboLock RNase inhibitor (Thermo Fisher Scientific) was added to the purified RNA to minimize degradation. All commercial kits were used according to the protocols provided by the manufacturer unless otherwise stated. Oligonucleotides used in this study can be found in **Table S1**.

### Samples and nucleic acid purification

Activated sludge and anaerobic digester biomass were obtained as frozen aliquots (−80°C) from the MiDAS collection (18). Sample metadata is provided in **Table S2**. Total nucleic acids were purified from 500 µL of sample thawed on ice using the PowerMicrobiome RNA isolation kit (MO BIO Laboratories) with the optional phenol-based lysis or with the RiboPure RNA purification kit for bacteria (Thermo Fisher Scientific). Purification was carried out according to the manufacturers’ recommendations, except that cell lysis was performed in a FastPrep-24 instrument for 4x 40 s at 6.0 m/s to increase the yield of nucleic acids from bacteria with sturdy cell walls (30). The samples were incubated on ice for 2 min between each bead beating to minimize heating due to friction. DNA-free total RNA was obtained by treating 41 µL of the purified nucleic acid with the DNase Max kit (MO BIO Laboratories), followed by clean up using 1.0x RNAClean XP beads with elution into 25 µL nuclease-free water.

### Primer-free full-length 16S rRNA library preparation and sequencing

Purified RNA obtained from biomass samples was pooled separately for each sample source type (activated sludge or anaerobic digester) to give equimolar amounts of 16S rRNA determined based on peak area in the TapeStation analysis software A.02.02 (SR1). Full-length SSU sequencing libraries were then prepared, as previously described (26). The SSU_rRNA_RT2 (activated sludge) and SSU_rRNA_RT3 (anaerobic digester) reverse transcription primers and the SSU_rRNA_l adaptor were used for the molecular tagging (**Table S1**), and approximately 1,000,000 tagged molecules from each pooled sample were used to create the clonal library. The final library was sequenced on a HiSeq2500 using on-board clustering and rapid run mode with a HiSeq PE Rapid Cluster Kit v2 (Illumina) and HiSeq Rapid SBS Kit v2, 265 cycles (Illumina), as previously described (26). Raw sequence reads were binned based on unique molecular tags, *de novo* assembled into synthetic long-read sequences, and trimmed equivalent to *E. coli* position 8 and 1507 using the fSSU-pipeline-RNA_v1.2.sh scripts script (https://github.com/KasperSkytte/AutoTax) (26).

### Primer-based full-length 16S rRNA gene library preparation and sequencing

The purified nucleic acids obtained from the biomass samples were pooled separately for each sample source type (activated sludge or anaerobic digester) with an equal weight of DNA from each sample. Full-length 16S rRNA sequencing libraries were then prepared, as previously described (26). The f16S_rDNA_pcr1_fw1 (activated sludge) or f16S_rDNA_pcr1_fw2 (anaerobic digester) and the f16S_rDNA_pcr1_rv (**Table S1**) were used for the molecular tagging, and approximately 1,000,000 tagged molecules from each pooled sample were used to create the clonal library. The final library was sequenced on a HiSeq2500 using on-board clustering and rapid run mode with a HiSeq PE Rapid Cluster Kit v2 (Illumina) and HiSeq Rapid SBS Kit v2, 265 cycles (Illumina) as previously described (26). Raw sequence reads were binned based on unique molecular tags, *de novo* assembled into synthetic long-read sequences, and trimmed equivalent to *E. coli* position 28 and 1491 using the fSSU-pipeline-DNA_v1.2.sh scripts script (https://github.com/KasperSkytte/AutoTax) (26).

### Extraction of high-quality “full-length” 16S rRNA gene sequences from SILVA

High-quality bacterial and archaeal 16S rRNA gene sequences in the SILVA 138 SSURef NR99 ARB-database was selected using the query pintail_slv = 100 and tax_slv = Bacteria* or tax_slv = Archaea*. Bacterial and Archaeal sequences were exported separately in the “fastawide” format after terminal trimming. Bacterial sequences were trimmed between the 27F and 1391R (31) primer binding sites equivalent to position 1,044 and 41,788 in the global SILVA alignment. Archaeal sequences were trimmed between the 20F (32) and the SSU1000ArR (33) primer binding sites equivalent to position 1,041 and 32,818 in the global SILVA alignment. A list of names for “full-length” sequences spanning the positions above was created using the Extract_full-length_16S_rRNA_names_from_SILVA.sh script, which takes advantage of the fact that ARB uses the period to specify terminal gaps and therefore indicates truncated sequences in the exported FASTA-files. The names were used to select and export the “full-length” bacterial or archaea sequences without trimming from the SILVA ARB-database.

### Generation of reference databases using AutoTax

AutoTax was created as a modular multi-step Linux BASH script that 1) generates FL-ASV and full-length 16S rRNA gene operational taxonomic units clustered at 99% identity (FL-OTU) reference sequences from high-quality, full-length 16S rRNA sequences. 2) Assigns a comprehensive seven-rank taxonomy to all reference sequences based on the SILVA taxonomy with the addition of placeholder names for unclassified taxa defined by *de novo* clustering of sequences using specific identity thresholds for each taxonomic rank (12). 3) Produces formatted reference databases, which can be directly used for classification using SINTAX or classifiers in the QIIME 2 framework.

AutoTax combines several software tools (GNU parallel v. 20161222 (34), USEARCH v. 11.0.667 (35), SINA v.1.6.0 (36), and R v.3.5.0 with the following packages: biostrings (37), doParallel (38), stringr (39), data.table (40), tidyr (41), and dplyr (42)) into a single BASH script that otherwise only requires a single FASTA file with the user provided full-length 16S rRNA gene sequences and the FASTA-formatted SILVA_138_SSURef_NR99_tax_silva reference database as input. The script, as well as a docker container image with all required software (except USEARCH, as the required 64 bit version it is non-free and must be purchased from https://drive5.com/usearch), is available on the GitHub repository https://github.com/KasperSkytte/AutoTax. The script is composed of separate, individual BASH functions to both allow for customization of the script as well as unit testing using the BASH Automated Testing System (https://github.com/bats-core/bats-core) where possible. Core functions of AutoTax are briefly described below. Expanded descriptions can be found in the supplementary information (**Text S1**).

#### Resolving full-length 16S rRNA amplicon sequence variant (FL-ASV)

Input sequences are oriented based on the SILVA 138 SSURef NR99 database using the *usearch - orient* command, dereplicated using *usearch -fastx_uniques* with the *-sizeout, -strand plus*, and *-threads 1* options, and finally denoised to produce the FL-ASVs using the *usearch -unoise3* command with the *-minsize 2* option.

#### Preparation of chimera-filtered full-length 16S rRNA OTUs (FL-OTUs)

Dereplicated sequences (before denoising) from above are clustered at 99% sequence identity using *usearch -cluster_smallmem* command with the *-id 0.99, -maxrejects 0, -centroids*, and *-sortedby size* options. Potential chimeras are identified and extracted using the *usearch -uchime2_ref* command with the *-strand plus, -mode sensitive*, and *-chimeras* options with the FL-ASVs from above as the reference database. The chimeras are finally removed to create the final FL-OTUs using the *usearch -search_exact* command with the *-strand plus* and *-dbnotmatched* options.

#### Taxonomy assignment

A complete taxonomy from kingdom to species is automatically assigned to each FL-ASV. In brief, the AutoTax script identifies the closest relative of each FL-ASV in the SILVA database using the *usearch -usearch_global* command, obtains the taxonomy for this sequence, and discards information at taxonomic ranks not supported by the sequence identity and the thresholds for taxonomic ranks proposed by Yarza *et al*. (12). The identity thresholds used for each of the taxonomic ranks are: Phylum (75.0%), Class (78.5%), Order (82.0%), Family (86.5%), Genus (94.5%), and Species (98.7%). For the species-level classification, the script identifies all type strains within the species-level threshold in the SILVA database and assigns a species-level classification to the FL-ASV if only a single species fits within the threshold. In addition, FL-ASVs are *de novo* clustered using the *usearch -cluster_smallmem* command using the thresholds for each taxonomic rank. The *de novo* clusters are labeled according to the format denovo_x_y, where x is a one-letter abbreviation for the taxonomic rank (k, p, c, o, f, g, and s), and y represents the FL-ASV number of the cluster centroid for each taxon. These labels act as a placeholder taxonomy, where the SILVA taxonomy does not provide any taxonomy information.

### Amplicon sequencing and analyses

Bacterial community analyses were performed by amplicon sequencing of the V1-3 variable region of the 16S rRNA gene as previously described (43) using the 27F (5’-AGAGTTTGATCCTGGCTCAG-3’) (31) and 534R (5’-ATTACCGCGGCTGCTGG-3’) (44) primers. Forward reads were processed using USEARCH v.11.0.667. Raw fastq files were filtered for phiX sequences using *usearch -filter_phix*, trimmed to 250 bp using *usearch -fastx_truncate - trunclen 250*, and quality filtered using *usearch -fastq_filter* with the *-fastq_maxee 1.0* option. The sequences were dereplicated using *usearch -fastx_uniques* with the *-sizeout* option. Exact amplicon sequence variants (ASVs) were resolved using *usearch -unoise3* (4). ASV-tables were created by mapping the raw reads to the ASVs using *usearch -otutab* with the *-zotus* and *-strand both* options. Taxonomy was assigned to ASVs using *usearch -sintax* with *-strand both* and *-sintax_cutoff 0.8* (13).

### Construction of phylogenetic trees and primer evaluation

FL-ASVs aligned to SILVA 138 NR99 ARB-database was obtained from the AutoTax output (temp/FL-ASVs_SILVA_aligned.fa) and loaded into SILVA 138 NR99 ARB-database. All bacterial FL-ASVs were selected and exported as a FASTA-file using the ssuref:bacteria positional variability by parsimony filter. A rough tree was created from the alignment using FastTree v. 2.1.10 (45) with the *-nt -gtr -gamma* options and loaded into ARB. The specificity of commonly used amplicon primers was determined for each FL-ASV using the *analyze_primers.py* script from Primer Prospector v. 1.0.1 (46). The specificity of primer sets was defined based on the overall weighted scores (OWS) for the primer with the highest score as follows: Perfect hit (OWS = 0), partial hit (OWS > 0, and ≤ 1), poor hit (OWS > 1). A comma-separated table with the specificity of each primer set for each FL-ASV was made in R, loaded into ARB, and used to color the tree.

### Data analyses and visualization

USEARCH v.11.0.667 was used for mapping sequences to references with *-usearch_global -id 0 - maxrejects 0 -maxaccepts 0 -top_hit_only -strand plus*, unless otherwise stated. Data were imported into R v.3.6.3 (47) using RStudio IDE (48), aggregated using the tidyverse package v.1.2.1 (https://www.tidyverse.org/), and analyzed and visualized using ggplot2 v.3.1.0 (49) and Ampvis2 v.2.4.0 (50).

### Data and code availability

Raw and assembled sequencing data is available at the European Nucleotide Archive (https://www.ebi.ac.uk/ena) under the project number PRJEB26558. The AutoTax script is available at https://github.com/KasperSkytte/AutoTax. AutoTax processed FL-ASV and FL-OTU expanded reference database in SINTAX and QIIME format, is available at https://figshare.com/articles/Data_used_in_AutoTax_paper/12377741/1. The AutoTax processed SILVA 138 SSURef NR99 database in SINTAX, and QIIME format is available at https://doi.org/10.6084/m9.figshare.12366626. R-markdown scripts used for data analyses and figures are available at https://github.com/msdueholm/Publications/tree/master/Dueholm2019a.

## Results and Discussion

### Sampling and high-throughput sequencing of full-length 16S rRNA sequences

To obtain 16S rRNA gene reference sequences for Danish wastewater treatment plants (WWTPs) and anaerobic digesters (ADs), we sampled biomass from 22 typical WWTPs and 16 ADs treating waste activated sludge located at Danish wastewater treatment facilities (**Table S2**). These facilities represent an important engineered ecosystem containing complex microbial communities of both bacteria and archaea, with the vast majority of microbes being uncultured and poorly characterized (51).

DNA and RNA were extracted and used for synthetic long-read 16S rRNA gene sequencing using both a primer-based and primer-free approach (26) (**Figure 1a**). A total of 926,507 full-length 16S rRNA gene sequences were obtained after quality filtering. These were denoised with UNOISE3 to generate a comprehensive reference database of 9,521 FL-ASVs with an error-rate below the detection limit according to theoretical calculations and analysis of 7,816 sequences previously obtained from the eight strain ZymoBIOMICS Microbial Community DNA Standard (26) (**Text S1**).

**Figure 1.**
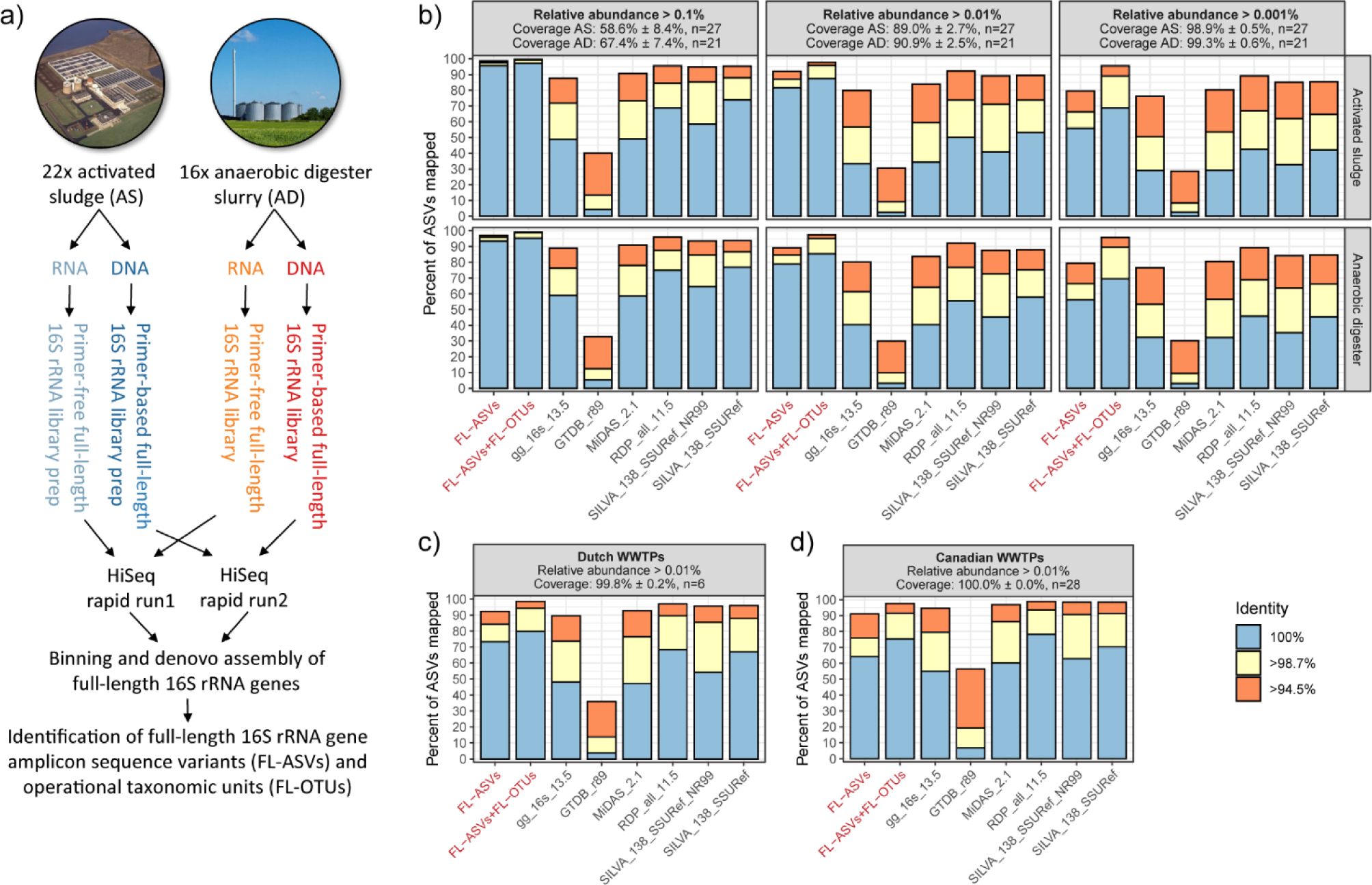
Construction and evaluation of the FL-ASV and FL-OTU expanded reference databases. a) Preparation of FL-ASVs and FL-OTUs. Samples were collected from WWTPs and ADs, and DNA and RNA were extracted. Purified DNA or RNA was used for the preparation of primer-based and “primer-free” full-length 16S rRNA libraries, respectively. These were sequenced and processed bioinformatically to produce the FL-ASVs and FL-OTUs b) Mapping of V1-3 amplicon data to the FL-ASV reference database, the FL-OTU expanded database, and commonly applied universal reference databases. ASVs were obtained from activated sludge and anaerobic digester samples and filtered based on their relative abundance before the analyses to uncover how well the databases cover the rare biosphere. The fraction of the microbial community represented by the remaining ASVs after the filtering (coverage) is shown as the mean ± standard deviation across plants. c) Mapping of V1-3 ASVs from Dutch WWTPs based on raw data from Gonzalez-Martinez et al. (57). For details, see Figure S2a. d) Mapping of V3-5 ASVs from Canadian WWTPs, based on raw data from Isazadeh et al. 2016 (56). For details, see Figure S2b.

To estimate the number of FL-ASVs belonging to novel taxa, FL-ASVs were mapped to the SILVA 138 SSURef NR99 database (15) using global mapping with USEARCH, and the identity of their closest relatives was compared to the thresholds for taxonomic ranks proposed by Yarza *et al.* 2014) (**Table 1**). The majority of the FL-ASVs (94%) had references in the SILVA database above the genus-level threshold (identity >94.5%), but 26% lacked references above the species-level threshold (identity > 98.7%), which are crucial for confident taxonomy assignment to ASVs.

**Table 1:** Numbers and percentage of FL-ASVs estimated to belong to novel taxa. FL-ASVs were mapped to SILVA 138 SSURef NR99 to find the identity with the closest relative in the database. The novelty was determined based on the identity for each FL-ASV using the taxonomic thresholds proposed by Yarza et al. (2014).

|  | Sequences | Percentage |
| --- | --- | --- |
| Phylum (<75.0%) | 1 | 0.01% |
| Class (<78.5%) | 1 | 0.01% |
| Order (<82.0%) | 3 | 0.03% |
| Family (<86.5%) | 16 | 0.17% |
| Genus (<94.5%) | 548 | 5.76% |
| Species (<98.7%) | 2449 | 25.7% |

### The FL-ASVs have better ecosystem coverage than universal reference databases

To evaluate if the FL-ASV database contained high-identity references for all bacteria in the ecosystem, we mapped V1-3 ASVs obtained from two sources: (1) the same samples used to create the FL-ASVs, and (2) samples from unrelated Danish WWTP and ADs. The ecosystem-specific FL-ASV database (9,521 seq.) included more high-identity references (>98.7% identity) for the abundant ASVs (relative abundance cutoff at 0.01%) in all samples analyzed, compared to the much larger universal databases: MiDAS 2.1 (548,447 seq.) (18), SILVA 138 SSURef NR99 (510,984 seq.) (15), SILVA 138 SSURef (2,225,272 seq.) (15), GreenGenes 16S v.13.5 (1,262,986 seq.) (14), and the full RDP v.11.5 (3,356,808 seq.) (16) (**Figure 1b and Figure S1**). ASVs were also mapped to the 16S rRNA database derived from the Genome Taxonomy Database (GTDB) release 89 (17,460 seq.) (52). However, this database lacked high-identity references for almost all ASVs. The poor coverage likely relates to the fact that 16S rRNA genes often fail to assemble in MAGs produced by short-read sequencing data (53). This problem will likely disappear in the future with the introduction of more high-quality MAGs with complete rRNA genes into the GTDB as a result of long-read sequencing technologies such as Nanopore and PacBio (54, 55).

When the rare biosphere was included in the analysis (relative abundance cutoff at 0.001%), a decrease in the percentage of ASVs with high-identity reference sequences was observed (**Figure 1b**). This may be a problem in ecosystems with high diversity, such as soil and sediments (9), where low-abundant microbes constitute a considerable fraction of the community, but also in engineered systems where transient or low abundant bacteria, such as pathogens or bacteria degrading micropollutants, may be important. To get a better representation of the rare biosphere, we created an additional reference database, which besides the FL-ASVs, contained the chimera-filtered, full-length 16S rRNA gene sequences clustered at 99% identity (FL-OTUs). This database greatly increased the coverage for the rare biosphere because it includes FL-OTUs that represent singletons. However, the improved coverage is achieved at the expense of taxonomic resolution (see later).

Since only Danish WWTPs and ADs were used to establish the comprehensive high-identity FL-ASV reference database, published amplicon data from non-Danish WWTPs (56, 57) was also evaluated (**Figure 1c-d, and S2**). Compared to the analyzed universal reference databases, the Danish reference FL-ASVs performed better or as well for most of the investigated non-Danish WWTPs, which indicates that even with less than 10,000 sequences, the database covers many of the microbes that are common to WWTP across the world, particularly for systems with nutrient removal. We anticipate that our ongoing sampling of more than 1,000 WWTPs and AD systems across all seven continents and different process-configurations (MiDAS Global, https://www.midasfieldguide.org/global) for high-throughput full-length 16S rRNA gene sequencing will provide references for the region-specific taxa in the near future, providing a comprehensive database of reference sequences for this ecosystem.

### A new comprehensive taxonomic framework

A major limitation in the classification of amplicon data from environmental samples is the lack of genus and species names for many uncultivated bacteria in the universal reference databases. To address this, we developed a simple taxonomic framework (AutoTax), which provides a consistent taxonomic classification of all reference sequences to all seven taxonomic ranks using identity thresholds (**Figure 2**).

**Figure 2.**
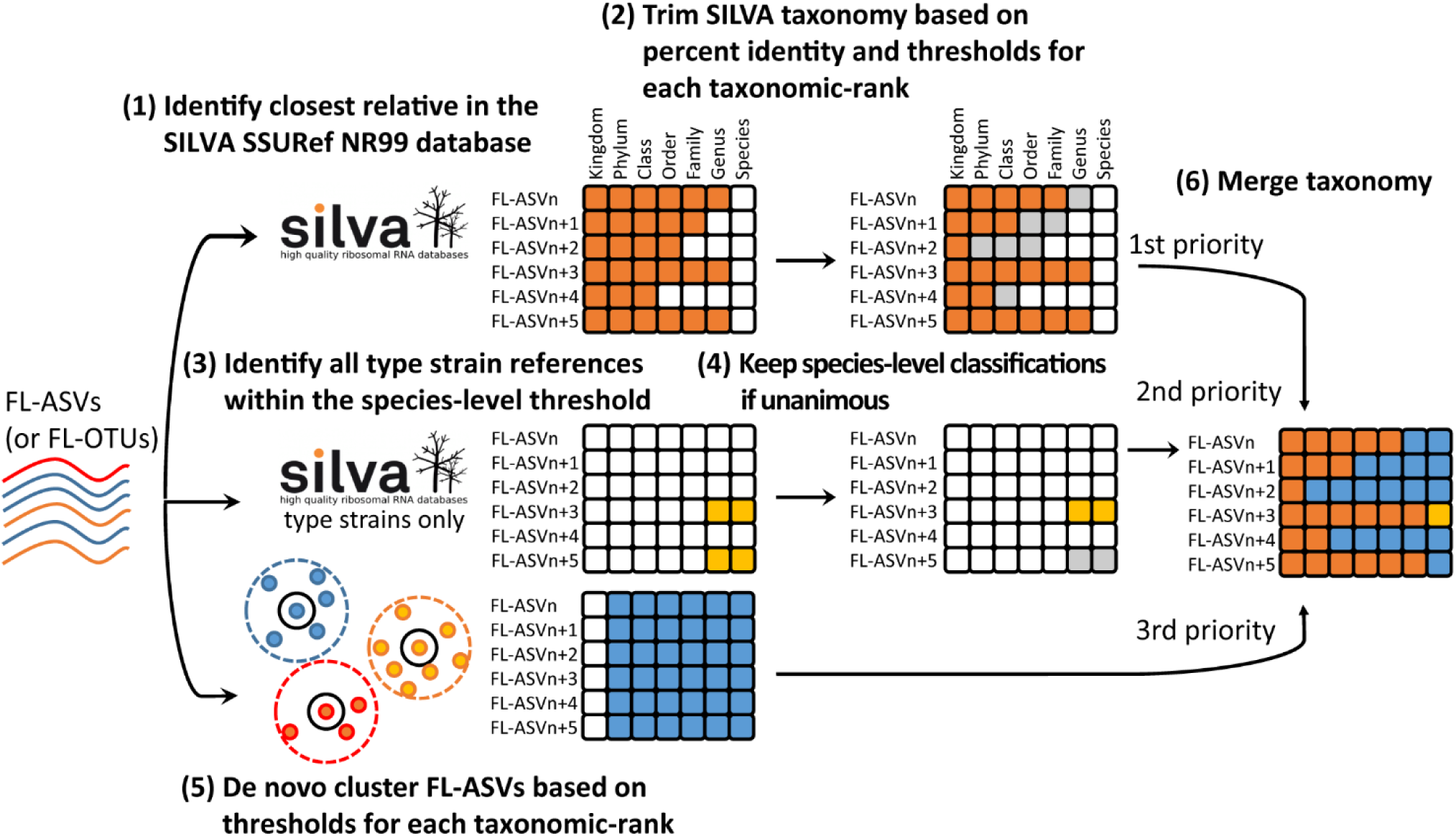
The AutoTax taxonomic framework. (1) FL-ASVs (or FL-OTUs) are first mapped to the SILVA 138 SSURef NR99 database to identify the closest relative and the shared percent identity. (2) Taxonomy is adopted from this sequence after trimming based on percent identity and the taxonomic thresholds proposed by Yarza *et al.* (12). To gain species-level information, FL-ASVs are also mapped to sequences from type strains extracted from the SILVA database, and species names are adopted if the identity is >98.7% and only a single species is within the threshold. (3) FL-ASVs are also clustered at different percent identity, corresponding to the thresholds proposed by Yarza *et al.* (12). The clustering is used to generate a stable *de novo* taxonomy. (4) Finally, a comprehensive taxonomy is obtained by filling gaps in the SILVA-based taxonomy with the *de novo*-taxonomy. Colored squares represent sources of taxonomic classifications of FL-ASVs: orange SILVA SSURef NR99, yellow: SILVA type strains, blue: *de novo* names, grey: names rejected during the AutoTax workflow.

In the AutoTax pipeline, the FL-ASVs (or FL-OTUs) are first mapped to the SILVA 138 SSURef NR99 database, which provides the taxonomy and percent identity of the closest relative in the database. This taxonomy is assigned to the FL-ASV down to the taxonomic rank supported by the sequence identity thresholds proposed by Yarza *et al.* (2014) (**Table 1**). Because of the stringent mapping and the taxonomy trimming based on percent identity, we obtained an overall better taxonomy assignment compared to commonly used classifiers, as revealed by a leave-one-out classification test (**Figure S3**).

Since species-level classification is desired wherever possible and the official SILVA taxonomy for bacteria and archaea is not curated at the species level (58), FL-ASVs were also mapped to the 16S rRNA gene sequences from type strains extracted from the SILVA 138 SSURef NR99 database as these carry official species names. Species-level classifications were assigned to the FL-ASVs if they shared more than 98.7% identity with only one species. If the FL-ASVs matched more than one, they were not classified at the species level due to the high risk of misclassification.

Because the SILVA taxonomy does not provide a complete seven-rank taxonomy for all sequences, the missing classifications are covered with a *de novo* placeholder taxonomy. This taxonomy was created based on the clustering of the FL-ASVs at identity thresholds corresponding to each taxonomic rank (12). The clusters were labeled according to the format denovo_x_y, where x is a one-letter abbreviation for the taxonomic rank (k, p, c, o, f, g, and s), and y represents the number of the FL-ASV which is the cluster centroid of the particular taxon. Because the applied clustering algorithm processes the sequences sequentially in the order they appear in the input file, sequences are always clustered in the same way, even if additional FL-ASVs are later added to the database. This strategy may not always yield the most optimal clusters, but the reproducibility is critical if the clusters are to be used as a robust placeholder taxonomy.

Merging of the SILVA- and the *de novo*-based taxonomies resulted in a few conflicts, e.g., where different FL-ASVs from the same species associate with more than one genus. In such cases, the genus-level classification for the centroid FL-ASV is adapted for all FL-ASVs within that species. However, these types of conflicts only applied to lower rank taxa (species), which were located close to the taxonomic threshold of the higher rank taxa (genus), and it only affected the classification of a low number of FL-ASVs (approx. 1%).

As AutoTax is based on the SILVA taxonomy, the taxonomy generated will change if another version of the SILVA SSURef NR99 reference database is used. Accordingly, users must specify which version of the SILVA database has been used when they publish databases created with AutoTax. We recommend that AutoTax generated databases are updated when there is a new version of the SILVA database. This ensures that the taxonomy is in agreement with the current central taxonomy.

### Taxonomy assignment to FL-ASVs with AutoTax

AutoTax provided placeholder names for many previously undescribed taxa in our FL-ASV database (**Table 2, Figure S4**). 95% of all species, 73% of all genera, 47% of all families, and 24% of all orders obtained placeholder names from the *de novo* taxonomy and would otherwise have remained unclassified. The novel taxa were affiliated with several phyla, especially the Proteobacteria, Planctomycetota, Patescibacteria, Firmicutes, Chloroflexi, Bacteroidota, Actinobacteriota, and Acidobacteriota (**Figure S4**). A prominent example is the Chloroflexi, where 5 out of 14 orders, 26 out of 34 families, and 142 out of 152 genera observed were assigned a *de novo* placeholder taxonomy. We believe that this will have important implications for future studies and the accumulation and transfer of knowledge about these taxa in WWTPs, given their high diversity and abundance, and their association with serious operational problems related to the settling of activated sludge (bulking) and foaming (59, 60). It should be noted that the placeholder taxonomy does not provide the same degree of support as traditional phylogenetic analyses, and should only be used until an official taxonomy is established.

**Table 2:** Numbers and percentage of taxa that were assigned *de novo* names.

|  | De novo taxa | Percentage |
| --- | --- | --- |
| Phylum | 1 | 2.22% |
| Class | 12 | 10.17% |
| Order | 66 | 24.35% |
| Family | 276 | 47.18% |
| Genus | 1324 | 72.91% |
| Species | 3973 | 95.28% |

### Improved classification of ASVs from WWTP and anaerobic digesters

To benchmark the FL-ASV database, we classified V1-3 amplicon data obtained from activated sludge and anaerobic digester samples (**Table S2**) using this database with the SINTAX-classifier and compared the results to classifications obtained using the universal reference databases (**Figure 3a**). With the use of the FL-ASV database, many more of the abundant ASVs (relative abundance >0.01%) were classified to the genus and species level (89.9±4.3% and 78.5±4.0%, respectively), compared to the use of popular public reference databases, including SILVA 138 SSURef NR99 (30.4±3.5% and 0%), GreenGenes 16s v. 13.5 (24.5±4.4% and 1.4±0.4%), GTDB r89 (22.1±2.8% and 11.6±1.2%), the RDP 16S v16 training set (20.3±4.0% and 0%), and even the MiDAS 2.1 (59.7±4.5% and 0.4±0.3%) - which is a manually ecosystem-specific curated version of the SILVA 123 SSURef NR99 database.

**Figure 3.**
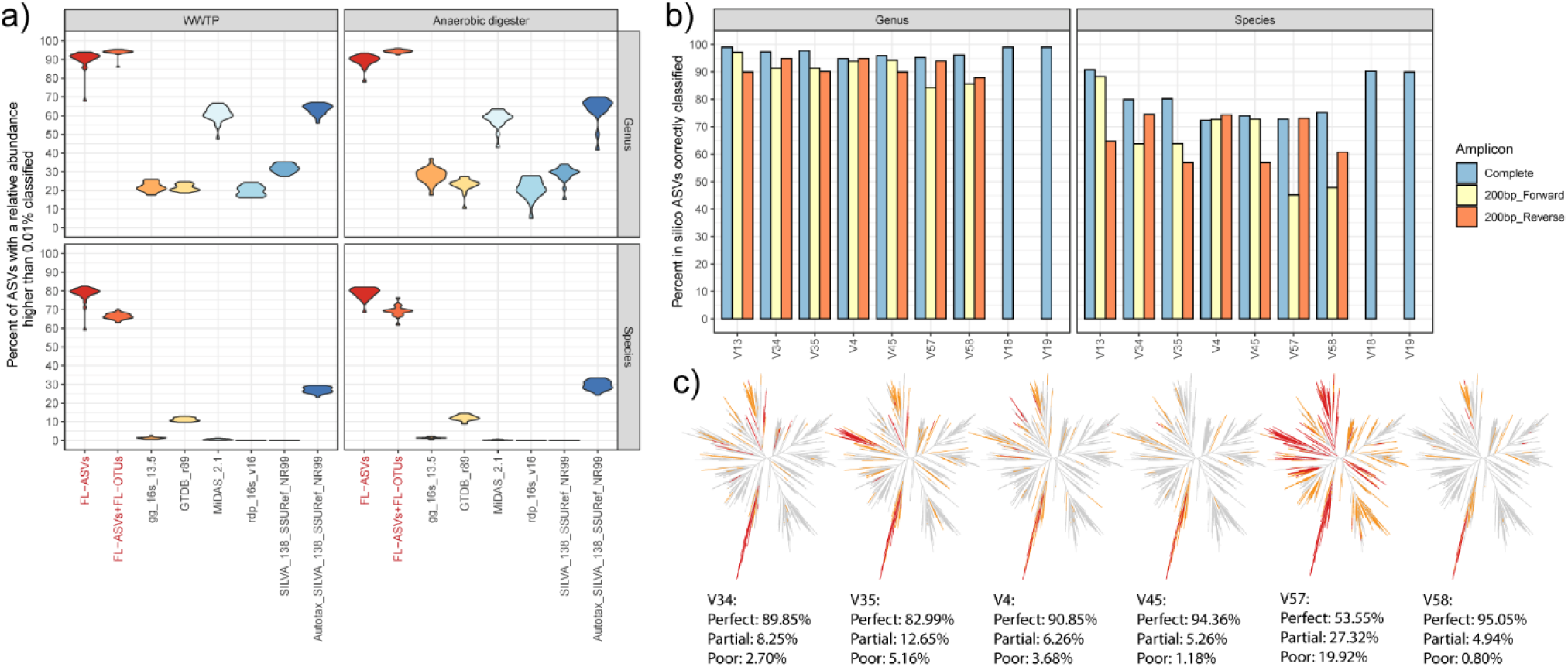
Evaluation of amplicon classification and primer specificity. a) The fraction of V1-3 ASVs from activated sludge and anaerobic digester sample with a relative abundance higher than 0.01% that were classified to the genus and species level using the FL-ASV reference database, the FL-OTU expanded database, and commonly applied universal reference databases, including the Autotax-processed SILVA 138 SSURef NR99 database. b) Classification of *in silico* bacterial ASVs, corresponding to amplicons produced using typical amplicon primers and the FL-ASV database. Results are shown for the complete amplicons as well as for partial amplicons corresponding to the first 200 bp from the forward or reverse read. c) Predicted primer bias associated with commonly used amplicon primers. The trees show the bacterial FL-ASVs, and the branch colors represent perfect hits (gray), partial hits (orange), and poor hits (red) (see materials and methods for definitions). The following primers were used: V13 (Lane 1991) (31), V18, V19, V34, and V58 (Klindworth et al. 2013) (62), V35 (Peterson et al. 2009) (64), V4 (Apprill et al. 2015) (65), V45 (Parada et al. 2016) (66), V57 (Chelius and Triplett 2001 and Bodenhausen et al. 2013) (67, 68).

We have here shown that increased coverage of the rare biosphere can be achieved by adding FL-OTUs clustered at 99% to the FL-ASV database. The FL-OTU expanded database increased the classification rate at the genus level for the abundant bacteria (relative abundance > 0.01%) (94.2±1.4% vs. 89.9±4.3%), but reduced classification at the species level (67.7±2.6% vs. 78.5±4.0%) (**Figure 3a**). When ASVs from the rare biosphere (relative abundance > 0.001%) were included in the analysis, the advantage of including FL-OTUs for genus-level classification was even more pronounced (93.0±0.6% vs. 80.7±1.7%), and an improved classification rate at the species level was also observed (69.9±1.5% vs. 66.8±2.4%) (**Figure S5**). We hypothesize that the reduced classification rate at the species level for the abundant ASVs is caused by sequencing errors as well as low-divergence chimera, which cannot be detected by chimera filtering (61). It should be noted that FL-OTUs are problematic for databases that are to be maintained and updated with additional references in the future. This is because the placeholder taxonomy will change if FL-OTUs are removed, and sequencing errors and chimeras are propagated if they are kept. We therefore recommend that FL-OTUs are only added for exploratory purposes.

To investigate the influence of AutoTax on taxonomic assignment, independent of our ecosystem-specific database, we applied the pipeline directly to high-quality, full-length sequences from the SILVA 138 SSURef_NR99 database. This increased the percentage of ASVs classified with SILVA at the genus level from 30.4±3.5% to 63.7±4.8%, and at the species-level from 0% to 28.2±2.4%, suggesting that the large universal databases would also benefit from the use of AutoTax. An advantage of the AutoTax processed SILVA database, which we have made publicly available, is that the placeholder taxonomy is universally applicable, providing a unique opportunity for studying the ecology of unclassified taxa across ecosystems.

### Taxonomic resolution of ASVs in combination with a comprehensive reference database

Classification of amplicon sequences can be challenging due to the limited taxonomic information in short-read sequences (12, 13). However, this may change with the access to reference databases with perfect references for the majority of all ASVs and a complete seven-rank taxonomy for all reference sequences. To determine the confidence of the amplicon classification in this scenario, we extracted ASVs *in silico* from the bacterial FL-ASVs corresponding to commonly amplified 16S rRNA regions, including full-length amplicons. These ASVs were classified against the FL-ASV database. We then calculated the fraction of amplicons correctly classified to the same genus and species as their corresponding FL-ASVs (**Figure 3b and Figure S6a**). Nearly all ASVs (95-99%) were assigned to the correct genus and most (72-91%) to the right species, depending on the taxonomic conservation of 16S rRNA region covered by the *in silico* amplicons. The primers targeting the V1-3 variable region performed exceptionally well for species-level identification (90.7% correct classifications), which is the same as for the full-length 16S rRNA gene amplicons. The commonly used primers targeting the V4 variable region were the worst (72.5% correct classifications). Very few of the sequences that did not receive the same taxonomic classification as their source reference sequence where misclassified (<0.2% at genus-level and <0.8% at species-level), with the majority not receiving any classification at the specific taxonomic rank (**Figure S6a**).

Sequencing costs on the Illumina platforms can be reduced considerably if single reads are used instead of merged reads. To evaluate the effect of reduced amplicon length, we compared the classification of 200 bp forward reads and reverse reads to those of full-length amplicons (**Figure 3b**). The decrease in the percentage of correct classifications was highly dependent on the 16S rRNA region targeted and from which direction the single reads were obtained. For the ecosystem studied here, the V1-3 and V4-5 forward read provided almost the same specificity as the full-length amplicons, whereas the reverse reads performed much worse for species-level classification. For the V3-4, V5-7, and V5-8, the reverse reads performed better than the forward reads, revealing the importance of choosing the right direction for single read amplicons.

To evaluate the effect of sequencing errors and low-divergence chimeras in the reference database, we also classified the *in silico* ASVs against the reference databases expanded with the FL-OTUs (**Figure S6b**). The result confirmed our prior observations that the inclusion of error prone references had a negative impact on our ability to classify short-read amplicons correctly; however, the effect was marginal at the genus-level. Full-length amplicons were less affected (**Figure S6b**), highlighting a clear advantage of using longer amplicons in combination with universal databases, which are likely to contain sequencing errors and low-divergence chimeras despite chimera filtering (61).

Overall, the analysis demonstrated that confident classification of short-read ASV sequences at the genus to species level is possible. However, it requires a reference database with a complete seven-rank taxonomy and perfect references for the majority of all ASVs. The scripts made available with this study can be used to confirm whether this is the case for samples from other studies.

### Ecosystem-specific evaluation of primer-bias

When choosing primers for amplicon sequence analyses, it is essential also to take primer-bias into account as some primer sets may result in ecosystem-relevant species being severely underestimated or absent from the analyses (30). The ecosystem-specific FL-ASV databases provide a near-perfect reference to determine the theoretical coverage of different primer sets for the given ecosystem so that an informed selection can be made (46, 62). It should be noted that primers used to generate the full-length 16S rRNA sequences for FL-ASV databases may introduce a bias, and we have here include primer-free (RNA-based) libraries to account for this. Evaluation of different primer sets using our FL-ASV database revealed a clear taxonomic bias associated with several primer sets (**Figure 3c**). The primers targeting the V4-5 and V5-8 regions had the best coverage of the FL-ASVs. The V5-7 primers demonstrated a very poor coverage. Because the FL-ASVs were trimmed after the forward priming site of the V1-3 primers, we were not able to evaluate the coverage of this primer pair here. However, it has previously been shown that the primers have a good overall agreement with metagenomic data for wastewater treatment systems and capture most of the process-critical organisms (31).

### Perspectives

High-throughput full-length 16S rRNA gene sequencing and automated taxonomy assignment (AutoTax) now allow individual research groups to develop their own FL-ASV ecosystem-specific reference databases for community profiling analyses. In addition, such databases can be used to evaluate the ecosystem-specific coverage and specificity of amplicon primers and fluorescence *in situ* hybridization (FISH) probes. The high quality of the FL-ASVs furthermore allows for the design of new primers and probes with improved confidence of the coverage and taxonomic resolution when applied within the target ecosystem. Collectively, the approach importantly allows for the identification and subsequent characterization of novel numerically important taxa for the specific environment, which would have otherwise been overlooked.

We acknowledge that the ability to generate new 16S rRNA gene reference databases quickly poses a risk for the development of several competing divergent taxonomies. We, therefore, recommend the custom databases are only used for exploratory purposes and combined with traditional phylogenetic analyses of key taxa. Ecosystem-specific databases that are broadly applied, such as the MiDAS database, should be created as open-source community efforts, and universal reference databases processed with AutoTax should be published and maintained in agreement with current developers of such databases. An important benefit of this approach is that the placeholder taxonomy can be used as a common language within the field, or in the case of universal reference databases, across all fields. This has major implications, e.g., in wastewater treatment systems, as it allows for the identification of unclassified taxa that are process-critical and decisive for process performance. If the current universal reference databases are used, the majority of ASVs will not get a genus-level classification, making it impossible to compare their prevalence across studies. Given that hundreds of amplicon-based studies are carried out every year worldwide, a considerable amount of useful information is lost when the data generated across studies is not comparable.

We have chosen to use the SINTAX-classifier for our analyses because it applies a simple algorithm that does not require training, and we expect the results to be less biased. However, some classifiers which use Bayesian inference, e.g., q2-feature-classifier (5) may yield better classifications. Kaehler et al. (63) recently demonstrated what environment-specific taxonomic abundance information could be used as weights for such classifiers to improve the accuracy of the taxonomy assignment. This approach is interesting because the abundance of raw full-length 16S rRNA gene sequences used to generate the FL-ASV database may be used as phylogenetically informative weights for the specific ecosystem.

We used the SILVA SSURef NR99 as the backbone taxonomy for AutoTax in this study, as this is currently the most comprehensive database. However, we anticipate the GTDB may replace SILVA in a future release of AutoTax when more high-quality genomes and MAGs with 16S rRNA genes (MIMAG standard (11)) are added to the database as the result of advances in long-read sequencing technology (Nanopore and PacBio). This will importantly link the 16S rRNA gene taxonomy with that derived from the more robust phylogenomic-based analyses, creating a unified language across the field of microbiology.

## Acknowledgments

We would like to thank the 22 wastewater treatment plants involved in the project for providing samples. This work was supported by the Danish Research Council [6111-00617A to P.H.N.]; and the Villum foundation [13351 to P.H.N.].

## Tables

**Table S1:**
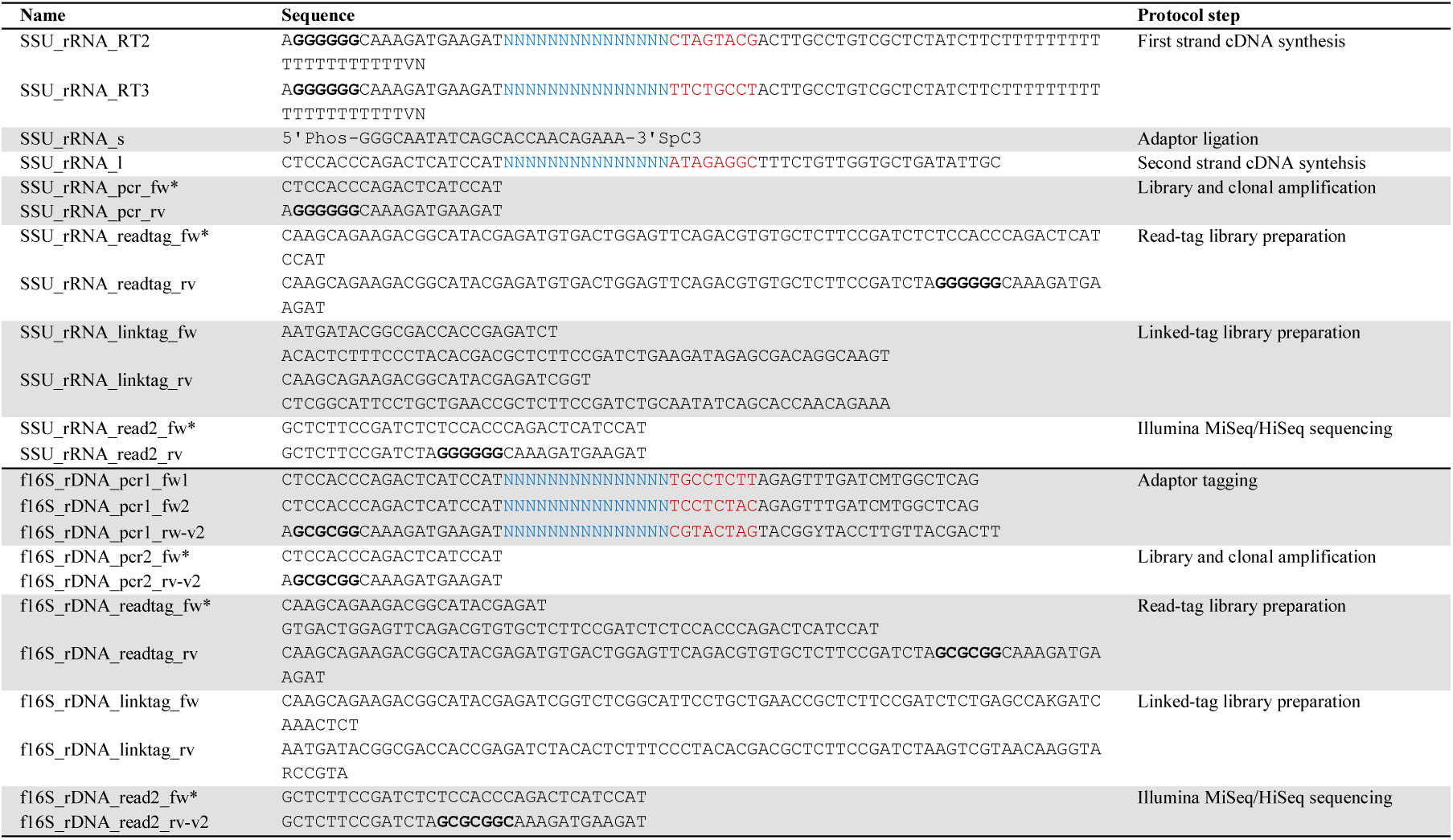
Oligonucleotides used for full-length 16S rRNA gene library preparation. Unique molecular tags, and sample barcodes are marked with blue and red, respectively. The GGGGGG motif marked with bold typefaces may form strong secondary structures known as guanine tetraplexes. This motif was replaced by GCGCGG in the DNA-based protocol, which increased the robustness of library preparation. Oligoes marked with “*” are identical between the RNA- and DNA-based protocols.

**Table S2:**
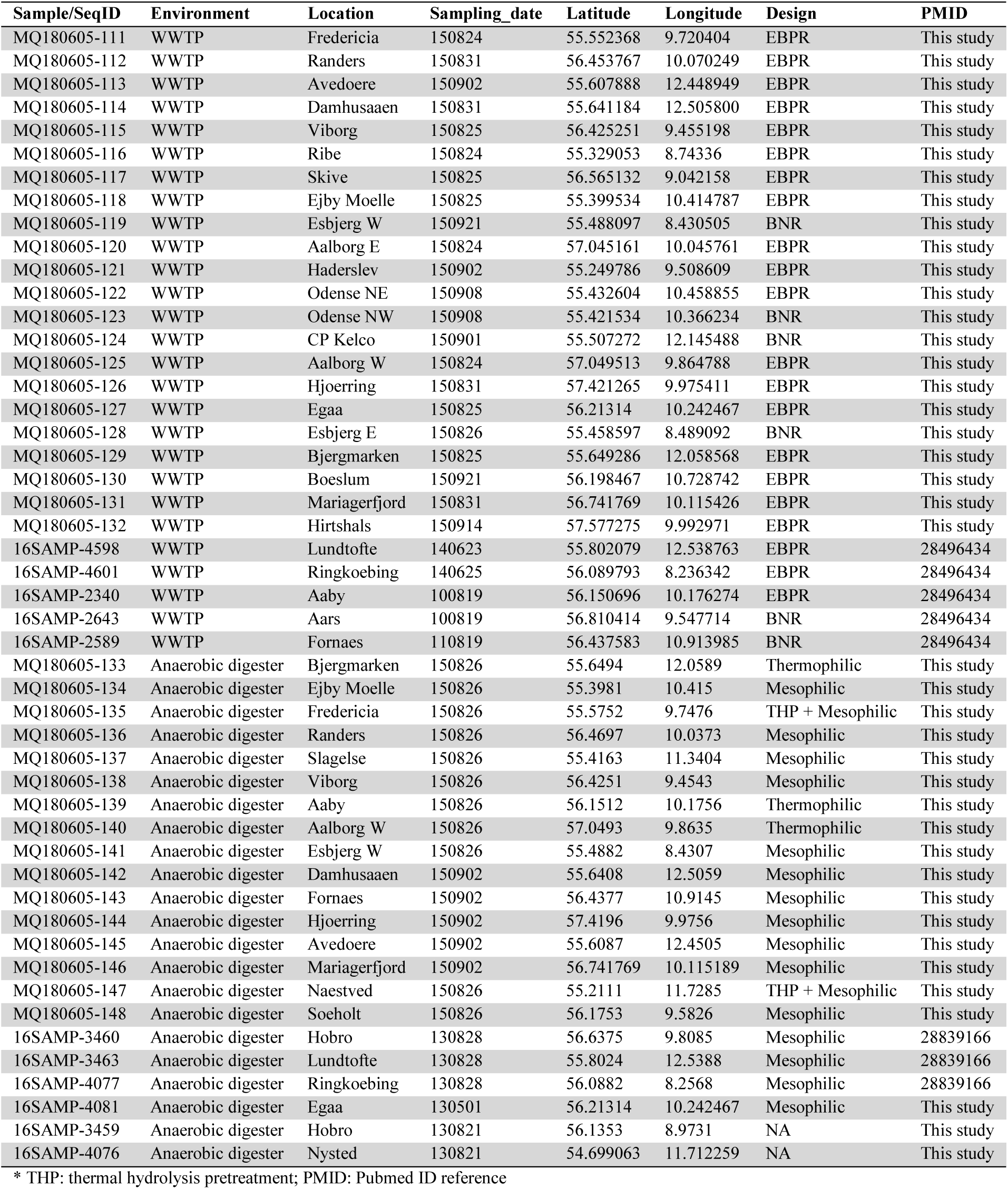
Samples used in this study.

| Sample/SeqID | Environment | Location | Sampling_date | Latitude | Longitude | Design | PMID |
| --- | --- | --- | --- | --- | --- | --- | --- |
| MQ180605-111 | WWTP | Fredericia | 150824 | 55.552368 | 9.720404 | EBPR | This study |
| MQ180605-112 | WWTP | Randers | 150831 | 56.453767 | 10.070249 | EBPR | This study |
| MQ180605-113 | WWTP | Avedoere | 150902 | 55.607888 | 12.448949 | EBPR | This study |
| MQ180605-114 | WWTP | Damhusaaen | 150831 | 55.641184 | 12.505800 | EBPR | This study |
| MQ180605-115 | WWTP | Viborg | 150825 | 56.425251 | 9.455198 | EBPR | This study |
| MQ180605-116 | WWTP | Ribe | 150824 | 55.329053 | 8.74336 | EBPR | This study |
| MQ180605-117 | WWTP | Skive | 150825 | 56.565132 | 9.042158 | EBPR | This study |
| MQ180605-118 | WWTP | Ejby Moelle | 150825 | 55.399534 | 10.414787 | EBPR | This study |
| MQ180605-119 | WWTP | Esbjerg W | 150921 | 55.488097 | 8.430505 | BNR | This study |
| MQ180605-120 | WWTP | Aalborg E | 150824 | 57.045161 | 10.045761 | EBPR | This study |
| MQ180605-121 | WWTP | Haderslev | 150902 | 55.249786 | 9.508609 | EBPR | This study |
| MQ180605-122 | WWTP | Odense NE | 150908 | 55.432604 | 10.458855 | EBPR | This study |
| MQ180605-123 | WWTP | Odense NW | 150908 | 55.421534 | 10.366234 | BNR | This study |
| MQ180605-124 | WWTP | CP Kelco | 150901 | 55.507272 | 12.145488 | BNR | This study |
| MQ180605-125 | WWTP | Aalborg W | 150824 | 57.049513 | 9.864788 | EBPR | This study |
| MQ180605-126 | WWTP | Hjoerring | 150831 | 57.421265 | 9.975411 | EBPR | This study |
| MQ180605-127 | WWTP | Egaa | 150825 | 56.21314 | 10.242467 | EBPR | This study |
| MQ180605-128 | WWTP | Esbjerg E | 150826 | 55.458597 | 8.489092 | BNR | This study |
| MQ180605-129 | WWTP | Bjergmarken | 150825 | 55.649286 | 12.058568 | EBPR | This study |
| MQ180605-130 | WWTP | Boeslum | 150921 | 56.198467 | 10.728742 | EBPR | This study |
| MQ180605-131 | WWTP | Mariagerfjord | 150831 | 56.741769 | 10.115426 | EBPR | This study |
| MQ180605-132 | WWTP | Hirtshals | 150914 | 57.577275 | 9.992971 | EBPR | This study |
| 16SAMP-4598 | WWTP | Lundtofte | 140623 | 55.802079 | 12.538763 | EBPR | 28496434 |
| 16SAMP-4601 | WWTP | Ringkoebing | 140625 | 56.089793 | 8.236342 | EBPR | 28496434 |
| 16SAMP-2340 | WWTP | Aaby | 100819 | 56.150696 | 10.176274 | EBPR | 28496434 |
| 16SAMP-2643 | WWTP | Aars | 100819 | 56.810414 | 9.547714 | BNR | 28496434 |
| 16SAMP-2589 | WWTP | Fornaes | 110819 | 56.437583 | 10.913985 | BNR | 28496434 |
| MQ180605-133 | Anaerobic digester | Bjergmarken | 150826 | 55.6494 | 12.0589 | Thermophilic | This study |
| MQ180605-134 | Anaerobic digester | Ejby Moelle | 150826 | 55.3981 | 10.415 | Mesophilic | This study |
| MQ180605-135 | Anaerobic digester | Fredericia | 150826 | 55.5752 | 9.7476 | THP + Mesophilic | This study |
| MQ180605-136 | Anaerobic digester | Randers | 150826 | 56.4697 | 10.0373 | Mesophilic | This study |
| MQ180605-137 | Anaerobic digester | Slagelse | 150826 | 55.4163 | 11.3404 | Mesophilic | This study |
| MQ180605-138 | Anaerobic digester | Viborg | 150826 | 56.4251 | 9.4543 | Mesophilic | This study |
| MQ180605-139 | Anaerobic digester | Aaby | 150826 | 56.1512 | 10.1756 | Thermophilic | This study |
| MQ180605-140 | Anaerobic digester | Aalborg W | 150826 | 57.0493 | 9.8635 | Thermophilic | This study |
| MQ180605-141 | Anaerobic digester | Esbjerg W | 150826 | 55.4882 | 8.4307 | Mesophilic | This study |
| MQ180605-142 | Anaerobic digester | Damhusaaen | 150902 | 55.6408 | 12.5059 | Mesophilic | This study |
| MQ180605-143 | Anaerobic digester | Fornaes | 150902 | 56.4377 | 10.9145 | Mesophilic | This study |
| MQ180605-144 | Anaerobic digester | Hjoerring | 150902 | 57.4196 | 9.9756 | Mesophilic | This study |
| MQ180605-145 | Anaerobic digester | Avedoere | 150902 | 55.6087 | 12.4505 | Mesophilic | This study |
| MQ180605-146 | Anaerobic digester | Mariagerfjord | 150902 | 56.741769 | 10.115189 | Mesophilic | This study |
| MQ180605-147 | Anaerobic digester | Naestved | 150826 | 55.2111 | 11.7285 | THP + Mesophilic | This study |
| MQ180605-148 | Anaerobic digester | Soeholt | 150826 | 56.1753 | 9.5826 | Mesophilic | This study |
| 16SAMP-3460 | Anaerobic digester | Hobro | 130828 | 56.6375 | 9.8085 | Mesophilic | 28839166 |
| 16SAMP-3463 | Anaerobic digester | Lundtofte | 130828 | 55.8024 | 12.5388 | Mesophilic | 28839166 |
| 16SAMP-4077 | Anaerobic digester | Ringkoebing | 130828 | 56.0882 | 8.2568 | Mesophilic | 28839166 |
| 16SAMP-4081 | Anaerobic digester | Egaa | 130501 | 56.21314 | 10.242467 | Mesophilic | This study |
| 16SAMP-3459 | Anaerobic digester | Hobro | 130821 | 56.1353 | 8.9731 | NA | This study |
| 16SAMP-4076 | Anaerobic digester | Nysted | 130821 | 54.699063 | 11.712259 | NA | This study |

\* THP: thermal hydrolysis pretreatment; PMID: Pubmed ID reference

**Table S3:**
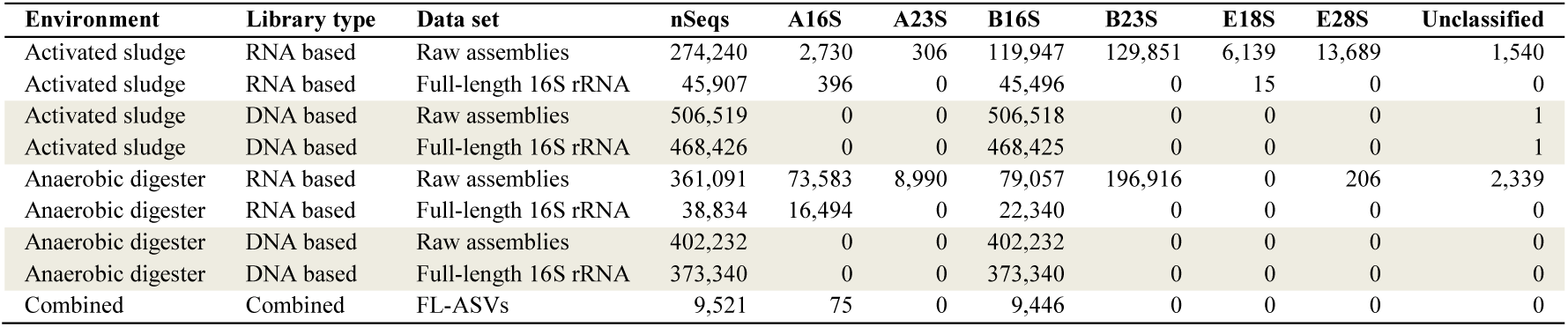
Classification of raw assemblies, full-length 16S rRNA, and FL-ASVs.

| Environment | Library type | Data set | nSeqs | A16S | A23S | B16S | B23S | E18S | E28S | Unclassified |
| --- | --- | --- | --- | --- | --- | --- | --- | --- | --- | --- |
| Activated sludge | RNA based | Raw assemblies | 274,240 | 2,730 | 306 | 119,947 | 129,851 | 6,139 | 13,689 | 1,540 |
| Activated sludge | RNA based | Full-length 16S rRNA | 45,907 | 396 | 0 | 45,496 | 0 | 15 | 0 | 0 |
| Activated sludge | DNA based | Raw assemblies | 506,519 | 0 | 0 | 506,518 | 0 | 0 | 0 | 1 |
| Activated sludge | DNA based | Full-length 16S rRNA | 468,426 | 0 | 0 | 468,425 | 0 | 0 | 0 | 1 |
| Anaerobic digester | RNA based | Raw assemblies | 361,091 | 73,583 | 8,990 | 79,057 | 196,916 | 0 | 206 | 2,339 |
| Anaerobic digester | RNA based | Full-length 16S rRNA | 38,834 | 16,494 | 0 | 22,340 | 0 | 0 | 0 | 0 |
| Anaerobic digester | DNA based | Raw assemblies | 402,232 | 0 | 0 | 402,232 | 0 | 0 | 0 | 0 |
| Anaerobic digester | DNA based | Full-length 16S rRNA | 373,340 | 0 | 0 | 373,340 | 0 | 0 | 0 | 0 |
| Combined | Combined | FL-ASVs | 9,521 | 75 | 0 | 9,446 | 0 | 0 | 0 | 0 |

## Figures

**Figure S1:**
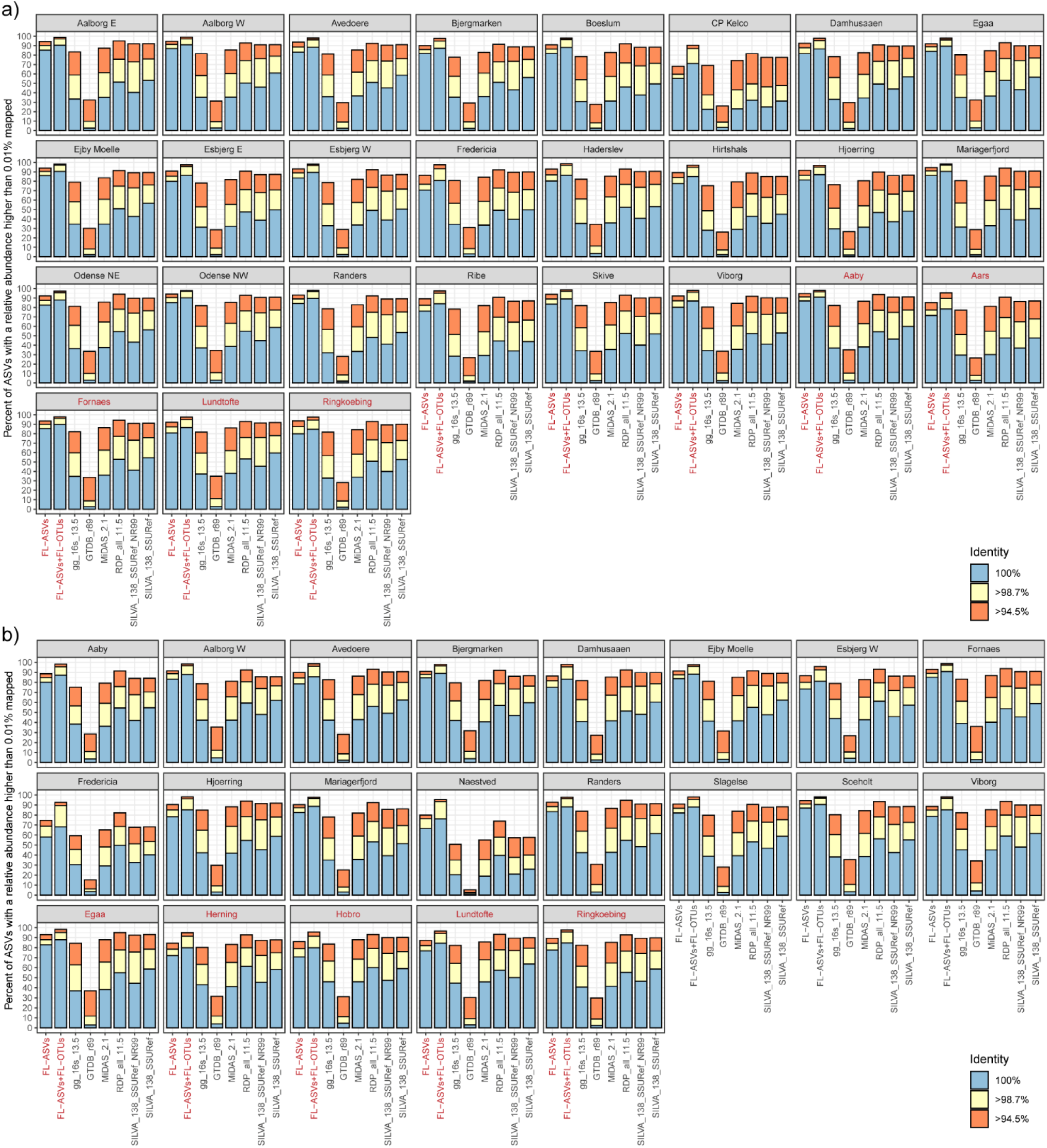
Mapping of activated sludge and anaerobic digester amplicon data to the FL-ASV and FL-OTU expanded reference database and public reference databases. V1-3 ASVs were obtained from the individual activated sludge (a) and anaerobic digester (b) samples used to create the FL-ASV reference database (black labels) and from other WWTPs (red labels).

**Figure S2:**
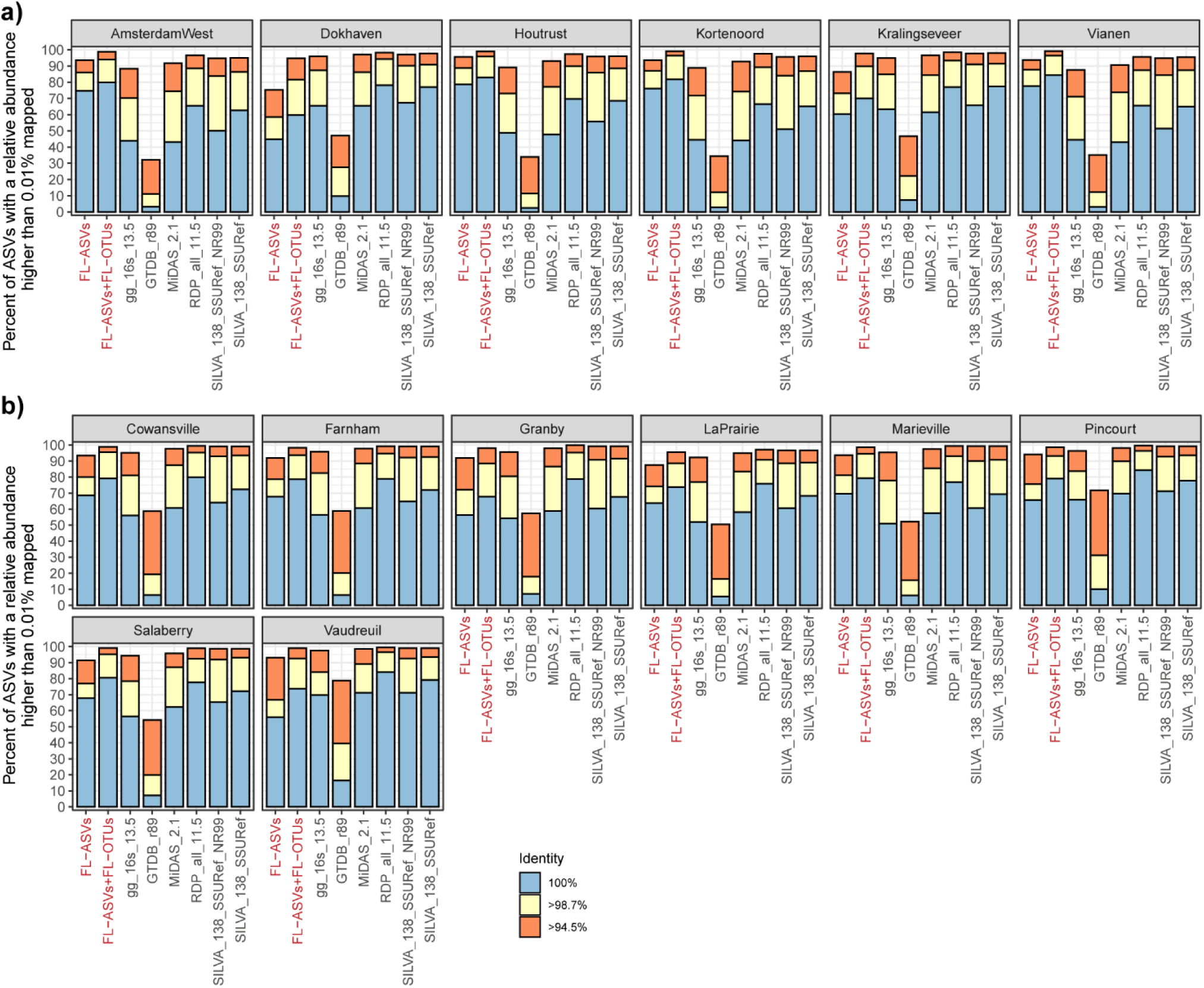
Mapping of amplicon data from non-Danish WWTP to the FL-ASV and FL-OTU expanded reference database and public reference databases. a) ASVs were created based on raw V1-3 amplicon reads obtained from Gonzarlez-Martinez *et al.* (57). Raw 454 genome sequencer reads in sff format were converted to fastq using CLC genomic workbench v. 20.0. Sequences were trimmed for the forward primer and truncated to 250 bp using *usearch - fastx_truncate -stripleft 19 -trunclen 250*, and quality filtered using *usearch -fastq_filter - fastq_maxee 1*. Processed reads were dereplicated using *usearch -fastx_uniques -sizeout*, denoised using *usearch -unoise3* (4). An ASV-table was produced with *usearch -otutab -zotus - sample_delim*. *-strand both*, and analyzed in R as described in the main article. All plants are based on conventional activated sludge except Dokhaven, which is configured with an Adsorption-Belebungsverfahren (AB) process. b) ASVs were created based on raw V3-5 amplicon reads obtained from Isazadeh *et al.* (2016)(56). Read 1 in SRA data were first oriented based on the SILVA 138 SSURef NR99 database using *usearch -orient*. Sequences were trimmed for the forward primer using *cutadapt -g CTACGGRNGGCWGC* (69), truncated to 250bp using *usearch - fastx_truncate -trunclen 250*, and quality filtered using *usearch -fastq_filter -fastq_maxee 1*. Processed reads were dereplicated using *usearch -fastx_uniques -sizeout*, denoised using *usearch - unoise3* (4). An ASV-table was produced with *usearch -otutab -zotus -sample_delim*. *-strand both*, and analyzed in R as described in the main article. The data originate from samples collected at eight Canadian WWTPs, which among others, differs in influent composition and treatment processes, for details see Isazadeh *et al.* (2016)(56). Marieville, Farnham, and Cowansville: Oxidation ditch. LaPrairie, Granby, Pincount, Salaberry: Conventional aeration. Vaudreuil: Sequencing batch reactor carrousel, which employs the principles of oxidation ditch.

**Figure S3.**
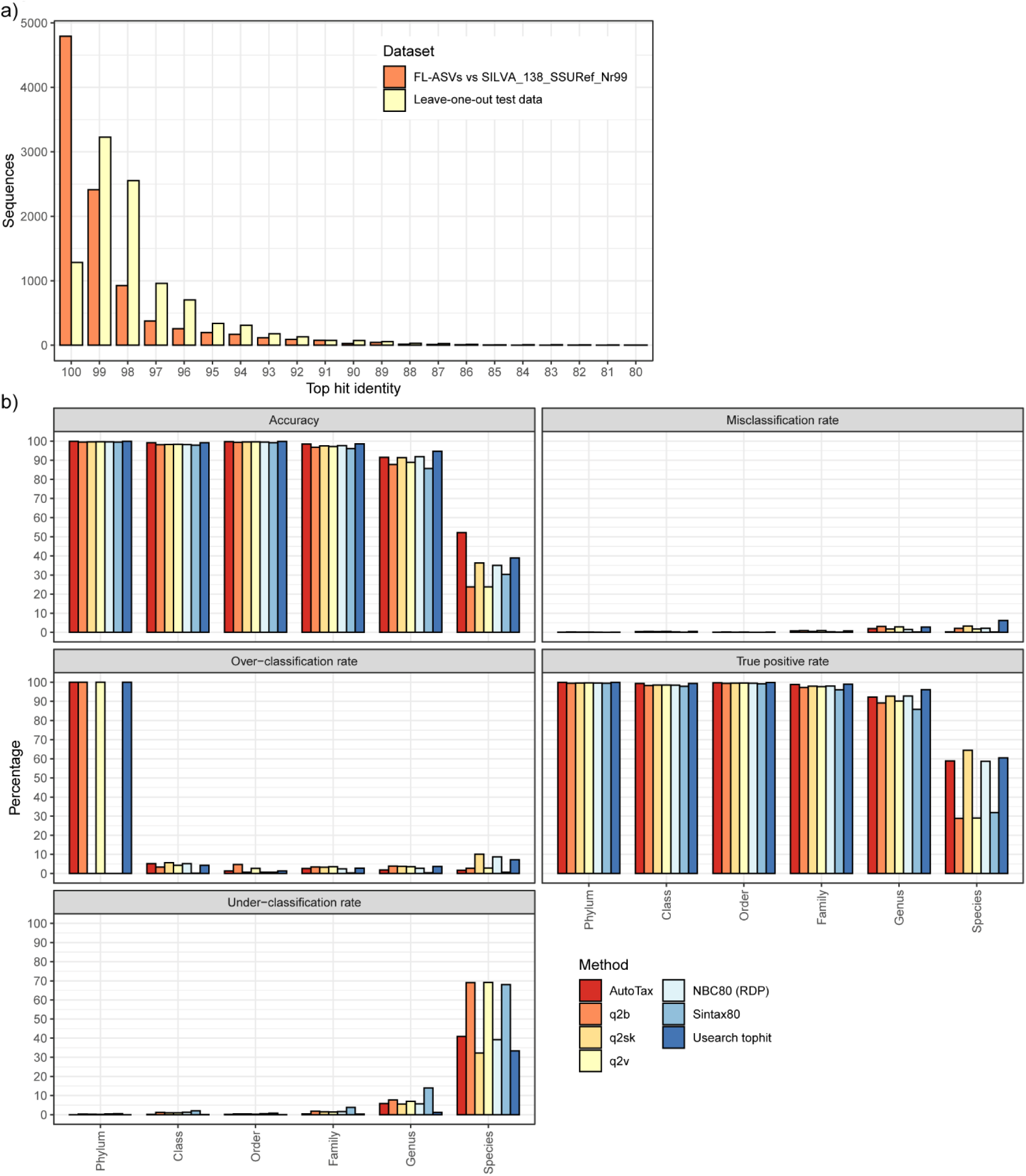
Evaluation of the taxonomy assignment to FL-ASVs. A leave-one-out validation test was performed using the AutoTax processed SILVA 138 SSURef NR99 database without the *de novo* taxonomy. This database only contains high-quality, full-length 16S rRNA sequences and improper-taxonomic entries (e.g., entries containing the words uncultured, unknown, incertae sedis, and possible) were removed. The database was divided into a 10,000 sequence test set, and the remaining database which was used for classification. a) Top hit identity distribution of the leave-one-out test data and for the FL-ASVs from this study mapped to the full SILVA 138 SSURef NR99 database (real data). The distributions are in the same range, confirming that the test data reflect actual data processed by AutoTax. b) Comparison of classification performance metrics for the AutoTax taxonomy assignment and commonly applied classifiers. The metrics used were the same as described by Edgar (13), but they were here applied to each taxonomic rank. Accuracy=TP/(K+OC); Misclassification rate=MC/K; Over-classification rate=OC/L; True positive rate=TP/K; Under-classification rate: UC/K, where K is the number of test sequences with a known taxonomy, L the number of test sequences without a known taxonomy, TP the number of names correctly assigned, MC the number of misclassification errors, OC the number overclassification errors, and UC the number of underclassification errors. The following classifiers were used: QIIME 2 feature-classifier with classify-consensus-blast (q2b), classify-consensus-vsearch (q2v), and classify-sklearn (q2sk) (5), USEARCH v.11 with SINTAX, NBC (Reimplementation of RDP), and USEARCH top hit (default parameters) (7, 13).

**Figure S4.**
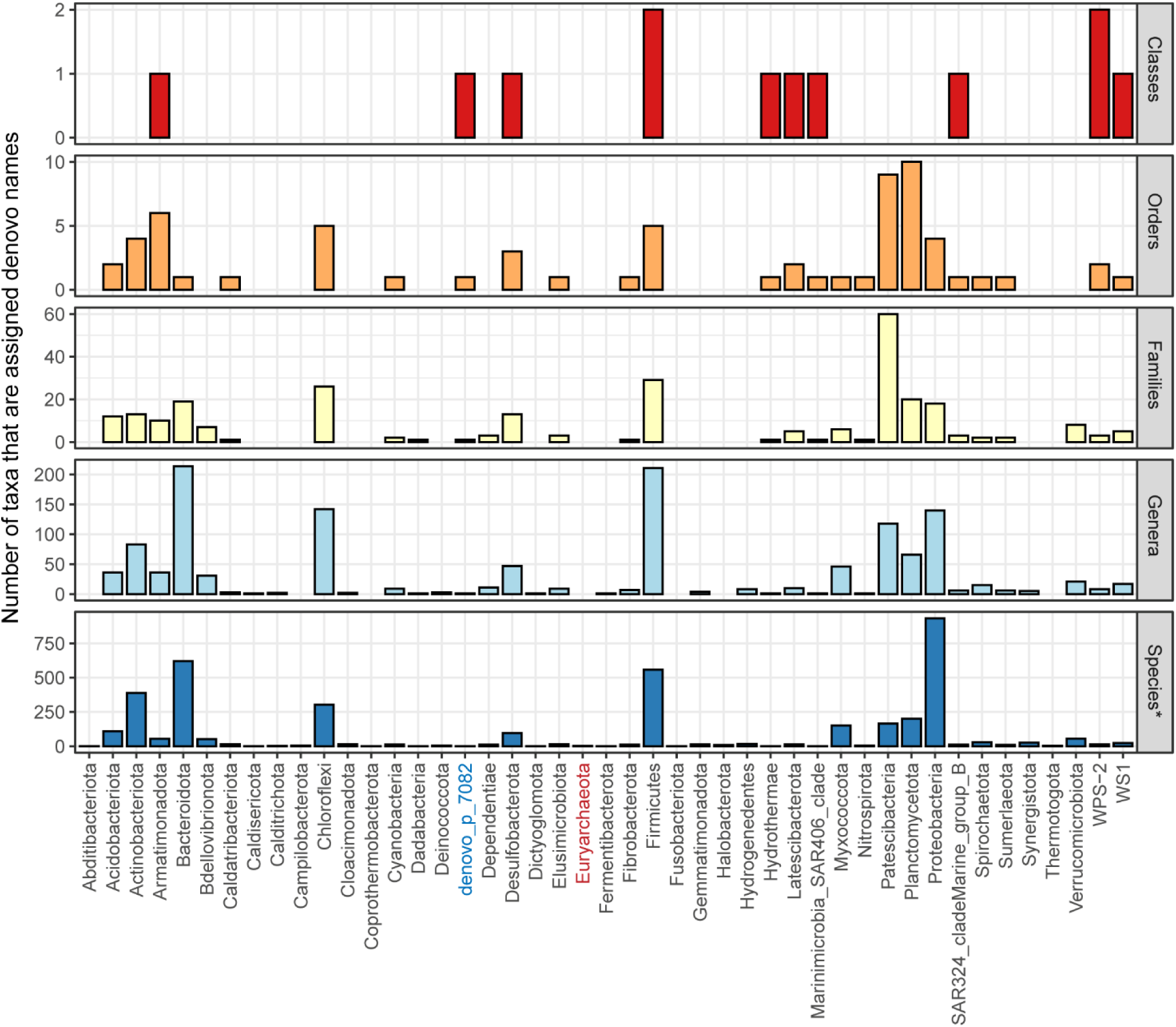
Novel taxa classified in the FL-ASV reference database. The number of taxa, which obtained names from the *de novo* taxonomy in different phyla. Archaeal phyla are highlighted in red and *de novo* phyla in blue.

**Figure S5.**
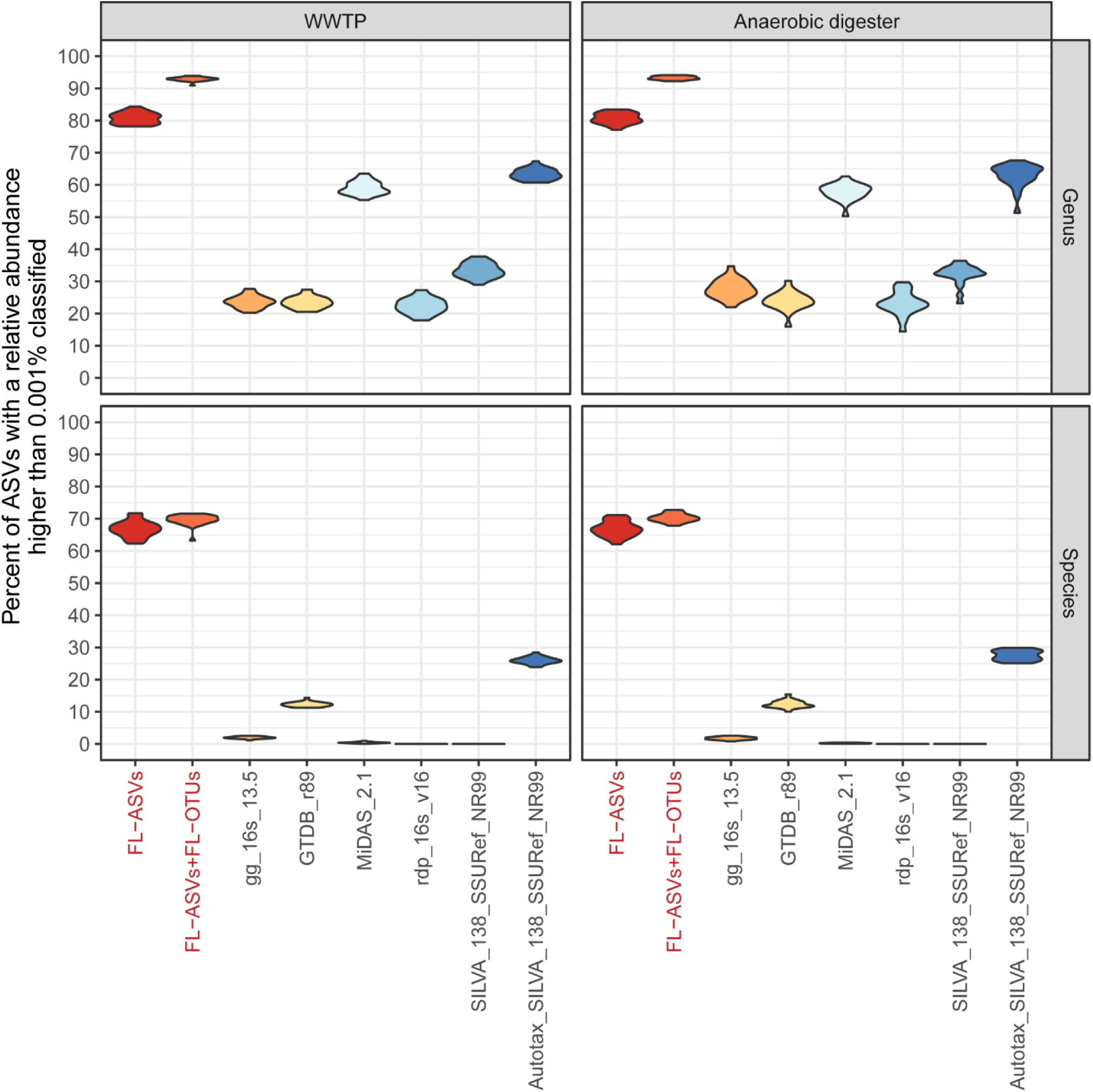
Effect of the rare biosphere on amplicon classification. a) The fraction of V1-3 ASVs from activated sludge and anaerobic digester sample with a relative abundance higher than 0.001% which was classified at the genus and species level using the FL-ASV reference database, the FL-OTU expanded database, and commonly applied universal reference databases, including the Autotax-processed SILVA 138 SSURef NR99 database.

**Figure S6.**
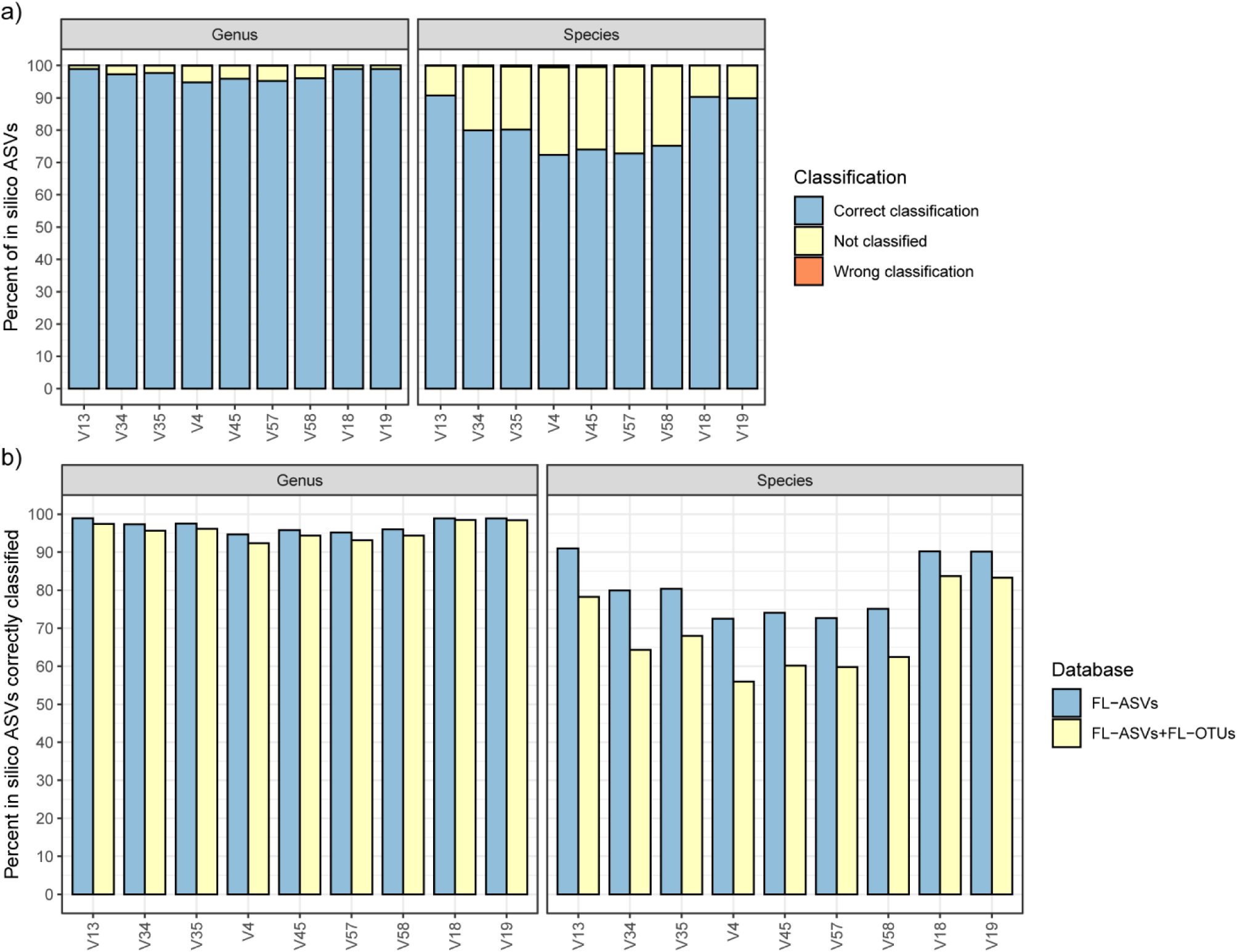
The specificity of taxonomy assignment using SINTAX and the FL-ASV and FL-OTU expanded reference database. a) Percentage of *in silico* bacterial ASVs, corresponding to amplicons produced using typical amplicon primers, that was correctly, wrongly, or not classified based on the FL-ASV database. b) Determining the effect of chimeras in the reference database. *In silico* bacterial ASVs were classified using the FL-ASV database and the FL-OTU expanded reference database.

## SUPPLEMENTARY INFORMATION FOR

### Detailed description of the AutoTax procedure

#### Full-length 16S rRNA exact amplicon sequence variant (FL-ASV) calling

Full-length 16S rRNA gene sequences are first oriented according to the SILVA 138 SSURef Nr99 database (1) using the *usearch -orient* command. The sequences are then dereplicated using *usearch -fastx_uniques* with the *-sizeout, -strand plus*, and *-threads 1* options. The *-sizeout* option put a size annotation (how many times a unique sequence was observed) in the FASTA header. The *-strand plus* option ensures that only correctly oriented sequences are considered when identifying identical sequences. The dereplicated sequences are sorted and numbered based on the size annotation. The *-threads 1* option ensures that sequences with the same size are always sorted and numbered the same way in the output. The dereplicated sequences are finally denoised to produce the FL-ASVs using the *usearch -unoise3* command with the *-minsize 2* option. The *-minsize* options specify the minimum abundance required for detecting an ASV. The default value for short-read amplicons is 8; however as the synthetic long-read sequences are independently amplified (due to the unique molecular identifiers (UMIs) added before the PCR steps) and have a very low error-rate we can lower the threshold to 2 with little risk of introducing sequences with errors, see “Calculation of FL-ASV error-rate” below.

#### Preparation of chimera-filtered full-length 16S rRNA OTUs (FL-OTUs)

Dereplicated sequences from above are clustered at 99% sequence identity to create FL-OTUs using *usearch -cluster_smallmem* command with the *-id 0.99, -maxrejects 0, -centroids*, and *-sortedby size* options. The *-maxrejects 0* option specified a complete database search, thereby providing more confident clusters. Potential chimeras are identified and extracted using the *usearch -uchime2_ref* command with the *-strand plus, -mode sensitive*, and -chimeras options using the FL-ASVs as a reference database. The sensitive mode was chosen as it captures more chimeras (on the expense of more false positives), as these are detrimental to the final reference database. The FL-ASVs were used as the reference database because it contains exact references for the abundant 16S rRNA genes, which are most likely to form chimeras. Chimeras are finally removed to create the chimera-filtered FL-OTUs using the *usearch -search_exact* command with the *-strand plus -dbnotmatched* options, using the identified chimera sequences as the query, and the pre-filtered FL-OTUs as the reference database.

#### Taxonomy assignment

For the taxonomy assignment, we first create two independent taxonomies. The first taxonomy is based on the most recent version of the SILVA SSURef Nr99 database and reflects the current state of microbial taxonomy. The second is a robust, although not necessarily evolutionary correct, *de novo* taxonomy. The latter will be used as a taxonomic placeholder for taxonomic ranks without information in the SILVA-based taxonomy.

The first step in both cases is the alignment of each FL-ASV to the global alignment in the SILVA reference database in ARB-format using SINA. The aligned sequences are hereafter trimmed to position 1048 to 41788 in the global SILVA alignment using the generic Linux command awk. The purpose of this trimming is two-fold. Firstly, it ensures that flanking sequences do not result in artificially low identities when the FL-ASVs are mapped to reference sequences in the SILVA database produced with the popular 27F and 1391R primers (2). Secondly, it allows a more robust *de novo* clustering (3). After trimming, gaps are removed from the alignment of FL-ASVs using the *usearch -fasta_stripgaps* command. Finally, the FL-ASVs are sorted based on the FL-ASV numbers in R.

The SILVA based taxonomy is obtained by mapping each of the trimmed FL-ASVs to the most recent version of the SILVA SSURef Nr99 and the type strain database in FASTA format and obtaining the taxonomic information from the closest relative as well as the percent identity. Mappings are done using the *usearch -usearch_global* command with the *-maxrejects 0, - maxaccepts 0, -top_hit_only, -strand plus, -id 0*, and *-blast6out* options for the complete SILVA database and the *-maxrejects 0, -maxaccepts 0, -strand plus, -id 0.987*, and *-blast6out* for the type strains. The *-maxrejects 0* and *-maxaccepts 0* options ensures that a comprehensive search is performed. The *-top_hits_only* and *-id 0* options provide the best hit in the complete SILVA database, whereas the *-id 0.987* without the *-top_hit_only* option provides all hits within the species-level threshold. The output files from the SILVA reference mapping are loaded into R, and a data frame with columns for FL-ASV number, percent identity, and SILVA taxonomy of the closest relative is created. The taxonomy field is then split into the seven main taxonomic ranks from kingdom to species. Fields that contain any of the following whole words are cleared: “uncultured”, “unknown”, “unidentified”, “incertae sedis”, “metagenome”, “bacterium”, and “possible”. In addition, we replace all “candidatus” with “Ca” and all white spaces with underscores. Finally, all characters except letters, numbers and period, dash, and underscore are removed.

The taxonomy obtained from the closest relative in the SILVA database does not necessarily match that of the FL-ASV. The taxonomy is therefore trimmed based on the percent identity between the FL-ASV and its closest relative. For this trimming we use the taxonomic thresholds proposed by Yarza *et al.* (4) (94.5% for genus-level, 86.5% for family-level, 82.0%, for order-level, 78.5% for class-level, and 75.0% for phylum-level). Species-level classifications are only obtained from the mapping against type strains, and no classifications are provided if reference sequences from more than one species are within the species-level threshold.

To generate a comprehensive *de novo* taxonomy, FL-ASVs are clustered based on the taxonomic thresholds proposed by Yarza *et al.* (4) using the *usearch -cluster_smallmem* command with the *-id x, -maxrejects 0, -uc*, and *-sortedby other* options. Where x represent the threshold for the given taxonomic rank. The cluster_smallmem algorithm with the *-sortedby other* option clusters the FL-ASVs based on when they appear in the input FASTA file, and the same clusters are therefore formed even though additional FL-ASVs are appended to the FL-ASV-database in the future. We confirmed this by processing only the first half of the FL-ASVs from this study, which provided identical clustering. The *-maxrejects 0* ensures that the best reference is found among all clustered sequences. The output file is a UCLUST-format tabbed text, which describes the clusters formed.

The six UCLUST output files (species- to phylum-level) are loaded into R, and each are converted into a data frame with two columns. The first column with clusters is named based on the taxonomic clustering rank, and the second column with the input sequences is named as the taxonomic rank below. Subsequently, the data frames are merged from species to phylum level. This results in a comprehensive taxonomy, where the clustered centroids determine affiliation to the taxonomic ranks above.

The SILVA-based taxonomy and the *de novo* taxonomy are finally merged in R by replacing empty fields in the SILVA-based taxonomy with the *de novo* taxonomic information. This results in a comprehensive taxonomy, where all FL-ASVs are described on all seven taxonomic ranks.

Merging of the two taxonomies results in a few cases where a taxon has more than one parent (e.g., sequences from the same species affiliate more than one genus). In these cases, the taxonomy of the FL-ASV with the lowest ASV-number within the taxa is assigned to all members. A log file is written with details of all such cases.

### Calculation of FL-ASV error-rate

The Unoise3 algorithm predicts which sequences represent true biological variants (ASVs) based on their relative abundance compared to those of closely related sequences with one or a few errors. Based on the error-rate profile of full-length 16S rRNA genes found in Karst et al. (5) and the assumption that sequencing-errors are randomly distributed we can calculate the relative chance of obtaining a correct ASV compared to an ASV with a single sequencing error when we only consider sequence seen at least twice:

The probability of obtaining a correct sequence for FL-ASV_n_ is the product of the probability of obtaining a sequence without any errors and relative abundance of FL-ASV_n_, and for the sequence to be seen twice, we need to square this number.

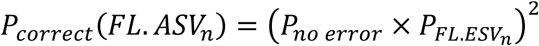

The probability of obtaining a sequence with a single error for FL-ASV_n_ is the product of the probability of obtaining a sequence with one error and relative abundance of FL-ASV_n_. However, to calculate the probability of picking a sequence with the same error, we need to divide this probability with three times the length of FL-ASV_n_ because the error could occur at any position in the sequence and could be any of the three wrong nucleotides. The probability for seeing a specific incorrect sequence twice therefore becomes:

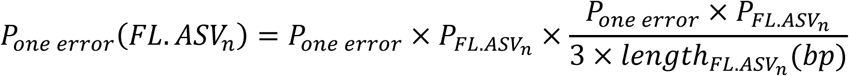

The relative chance of obtaining a correct FL-ASV compared to an FL-ASV with a single sequencing error is therefore:

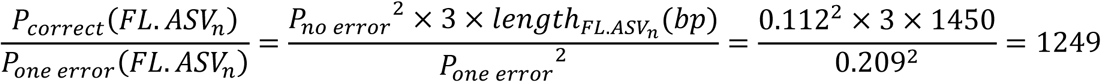

The results show that for each time we obtain an FL-ASV with a single error, we should expect approximately 1250 times more sequences of the correct FL-ASV. The difference should be large enough for Unoise3 to be able to remove almost all sequences with errors even though the errors are not entirely randomly distributed.

To confirm this, we reanalyzed the mock data used in Karst *et al.* (5). This data contained 7816 trimmed sequences (27f to 1492r) from a balanced mixture of DNA from *Escherichia coli* MG1655 (NC_000913), *Bacillus subtilis* subsp. subtilis str. 168 (NC_000964) and *P. aeruginosa* PAO1 (NC_002516). Denoising with *usearch -unoise3* command and the *-minsize 2* option produced 16 error-corrected sequences, of which 13 mapped perfectly to the 16S rRNA reference sequences, two mapped with a single error, and one mapped with five errors. As the two FL-ASVs with single errors were observed 14 and 4 times, respectively, it is highly unlikely that they represent sequencing error based on the previous error-rate calculations. The same is holds for the FL-ASV with five errors, although this sequence was observed only twice.

